# Pathological tau leads to network hyperexcitability through neuronal-subtype-specific functional impairment

**DOI:** 10.64898/2026.10.05.756807

**Authors:** Changyi Ji, Yunlu Zhu, Hediye Erdjument-Bromage, Soomin C Song, Natalie Chang, Amanda R Liu, Alexa R Lee, Olivia DeGulis, David Schoppik, Thomas A Neubert, Hyung Don Ryoo, Einar M Sigurdsson

**Author notes:** Corresponding author: Changyi Ji, Department of Neuroscience, Institute for Translational Neuroscience, New York University Grossman School of Medicine Science Building, 11th floor, 435 East 30th Street New York, NY 10016, Einar M. Sigurdsson Department of Neuroscience Department of Psychiatry, Institute for Translational Neuroscience, New York University Grossman School of Medicine Science Building, 11th floor, 435 East 30th Street New York, NY 10016.

## Abstract

Pathological tau disrupts neuronal activity and brain function in Alzheimer’s disease and other tauopathies. However, the underlying mechanisms for neuronal network dysfunction remain largely unknown. Using human inducible pluripotent stem cell (iPSC) - derived neurons, we show that the familial *MAPT* P301L mutation leads to neuronal network hyperexcitability. Imbalanced excitation and inhibition were attributable to neuronal subtype-specific tau pathology that was associated with different alterations of tau interactomes in excitatory versus inhibitory neurons. Suppressing enhanced excitatory synaptic transmission or blocking Ca^2+^ influx resulting from enhanced P301L tau/AHNAK interaction in inhibitory neurons rescued the neuronal network hyperexcitability. This study expands our understanding on how disease-causing tau mutations alter the excitatory/inhibitory balance, leading to network dysfunction in human neurons.

**HIGHLIGHTS:**

- The FTD-causing P301L tau mutation leads to neuronal network hyperexcitability.
- P301L tau causes hyperactivity of excitatory neurons while hypoactivity of inhibitory neurons.
- Tau and P301L tau interactomes differ in excitatory vs. inhibitory neurons.
- Enhanced excitatory synaptic transmission and P301L tau/AHNAK interaction in inhibitory neurons contribute to neuronal network hyperexcitability.

**Graphic Summary:** 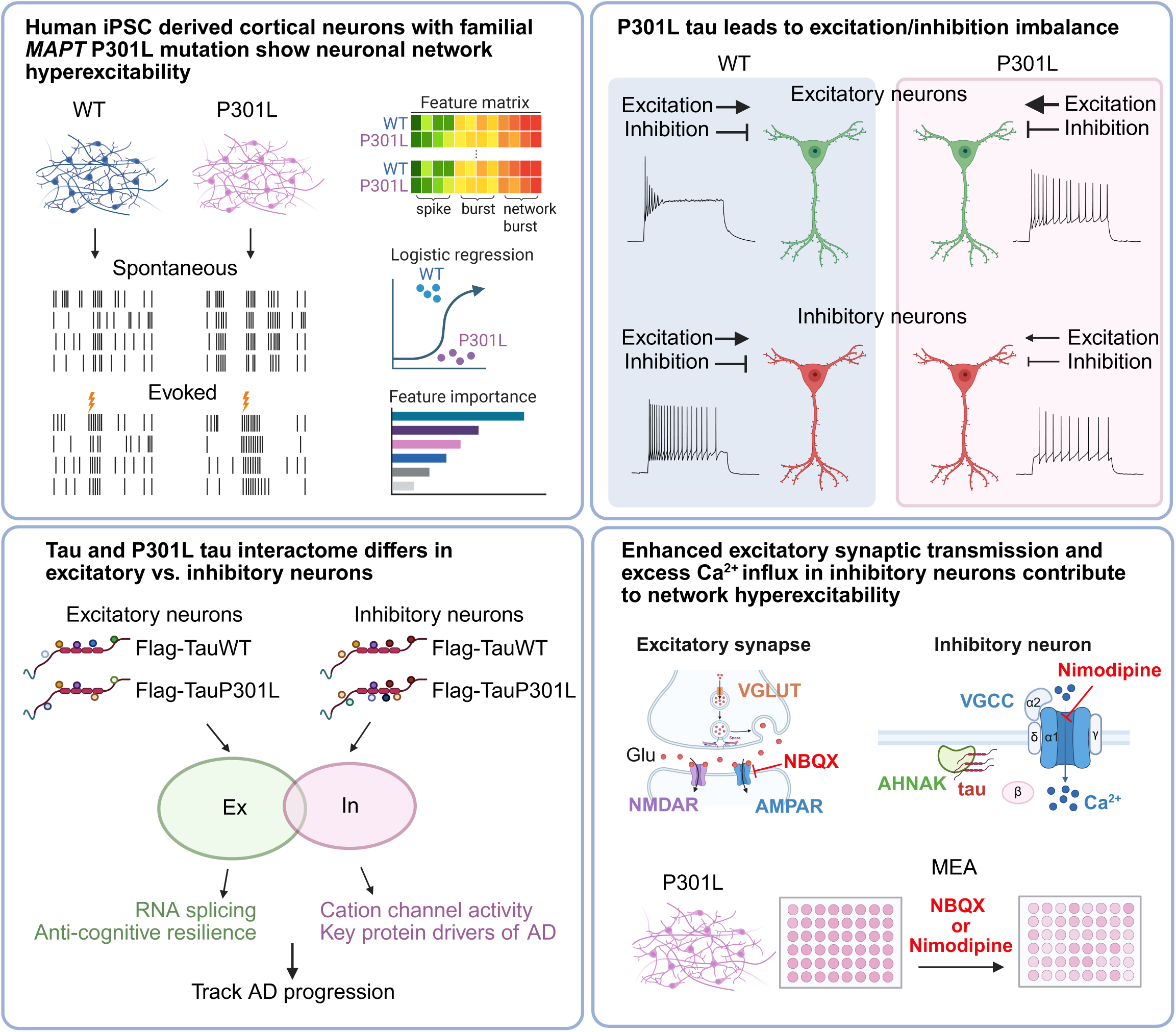

## INTRODUCTION

The accumulation of pathological tau is a central hallmark of Alzheimer’s disease (AD) and frontotemporal dementia (FTD) associated with tauopathy, which is closely correlated with cognitive decline and other behavioral deficits in patients [1]. Under physiological conditions, tau is associated with microtubules and plays important roles in axonal transport and vesicle trafficking [2,3]. During disease, pathological tau fails to stabilize microtubules and compromises neuronal function [2,3]. Emerging evidence suggests that certain neuronal subtypes are selectively vulnerable to tau pathology [4–8]. Our previous study of JNPL3 tauopathy mice, which carry human 0N4R tau P301L mutation, observed selective loss of GABAergic interneurons in the hippocampus [4]. Other studies suggested the loss of glutamatergic neurons in the entorhinal cortex [5,8] and impairment of glutamatergic excitation [7], which contributed to spatial memory deficits in a tauopathy mouse model [5,6]. These diverse observations allude to the complex, spatial and temporal characteristics of tau pathogenesis. Synaptic and neural network dysfunction are widely observed as early pathological events before neurodegeneration in tauopathy [9–11]. However, few studies have examined the mechanisms underlying the impact of pathological tau on neuronal and network dysfunction, which are likely attributable to selective cell loss and/or altered neuronal activity in different neuronal subtypes depending on disease stage.

Specifically, there has been limited research on the functional impact of tau on different neuronal subtypes. In transgenic *MAPT* knockout mice, global genetic ablation of tau affected the activity of excitatory and inhibitory neurons differently [12]. While excitatory neurons showed reduced spontaneous firing, inhibitory neurons exhibited enhanced excitability and reduced homeostatic responses to stimulation [12]. Reduced excitation over inhibition counteracted network hypersynchrony in primary mouse neurons, suggesting different physiological roles of tau in neuronal subtype function [12]. Our recent study on JNPL3 tauopathy mice overexpressing human mutant P301L tau also observed differential functional impact of pathological tau on fast-spiking (FS) interneurons, presumably inhibitory neurons vs. non-fast-spiking (non-FS), presumably excitatory neurons [13]. While the presence of mutant tau significantly increased the excitability of both, the extent of influence was larger in non-FS neurons [13]. Moreover, the excitatory synaptic transmission was significantly enhanced in FS neurons whereas non-FS neurons received less excitation [13]. These imbalanced changes were accompanied by hypersynchrony of the local motor cortical neuronal circuit, as well as impaired responses of neuronal ensembles in running tauopathy mice [13]. Taken together, these two studies suggested cell-subtype specific roles of tau at both physiological and pathological levels. The molecular and cellular mechanisms underlying differential influences of pathological tau on neuronal subtypes warrant further investigation.

Patient-derived iPSC-differentiated neurons (iPSC-iN) are a valuable disease relevant model, which provides unique insights into human disease etiology and pathogenesis. Previous studies using iPSC-iNs with familial tau mutations demonstrated that these mutations are sufficient to induce global transcriptomic changes [7,14,15]. Neurons carrying the *MAPT* R406W mutation exhibited decreased expression of various GABA receptors, suggesting impaired post-synaptic GABAergic signaling [14,16]. Familial tau mutations also alter neuronal activity. Human neurons carrying *MAPT* IVS10+16 mutation with enhanced 4-repeat (4R) over 3-repeat (3R) tau expression exhibited impaired excitability [17,18]. However, the mutant neurons received increased excitatory synaptic transmission and enhanced Ca^2+^ influx, mediated by mitochondrial reactive oxygen species [17,19]. Additionally, mutant V337M tau impaired axon initial segment plasticity and altered excitability homeostasis in excitatory neurons [20]. Furthermore, a selective loss of glutamatergic neurons was observed in long-term culture of iPSC-derived brain organoids with tau V337M mutation [7]. While the pathological influences of familial *MAPT* mutations vary, the functional impact of the same tau mutation among different neuronal subtypes has not been directly compared side by side in human neuronal models. The differential changes in neuronal subtypes likely contribute to the neuronal network dysfunction in tauopathy.

To understand the complex interplay between selective neuronal vulnerability and neuronal network dysfunction in tauopathy, we investigated whether tau pathology affects the function of different neuronal subtypes as well as neuronal network activity in iPSC-iN carrying *MAPT* P301L mutation. This mutation, located in exon 10 of 4R tau, causes FTD with atrophy of the frontal and temporal lobes, basal ganglia and hippocampus. In iPSC-iNs, neuronal networks with P301L mutant tau displayed hyperactivity, showing less homeostatic regulation in response to repeated electrical stimulation. Altered cellular composition was observed in these iPSC-iNs and the mutant neurons accumulated phospho-tau in inhibitory neuronal soma, as well as excitatory and inhibitory synapses. Electrophysiological studies found that mutant tau differentially affected firing properties and synaptic transmission of excitatory and inhibitory neurons. To further investigate the molecular and cellular mechanisms, we profiled specific tau interactomes in excitatory vs inhibitory neurons by immunoprecipitation followed by mass spectrometry. We have identified novel tau interactors in these neuronal subtypes, some of which were represented in human brain protein modules associated with AD or key driver proteins in an anti-cognitive resilience protein network. Following validation of a candidate tau interactor AHNAK and various pharmacological inhibition experiments, the neuronal hyperexcitability and abnormal network activity could be reversed by blocking the enhanced excitatory synaptic transmission, or Ca^2+^ influx through the L-type voltage gated Ca^2+^ channel in inhibitory neurons. Furthermore, overexpression of mutant tau in a healthy subject background confirmed the dominant influence of mutant tau expressed in inhibitory neurons on their neighboring excitatory neurons and led to neuronal network hyperexcitability. Lastly, we also extracted electrophysiological signatures from these neuronal network recordings and trained a logistic regression model that could robustly classify functional phenotypes of iPSC-iNs carrying P301L mutations from isogenic controls. Taken together, mutant tau in inhibitory neurons plays an influential role in cortical neuronal network dysfunction at an early stage of tauopathy. Our study reveals the differential functional impact of *MAPT* P301L mutation on neuronal subtypes and provides a mechanistic insight into early tau pathogenesis that can be targeted for therapy.

## RESULTS

### Neuronal and network hyperactivity in neurons with pathological tau

Patient derived iPSCs carrying *MAPT* P301L mutation and their isogenic control were differentiated into forebrain cortical neurons (Fig. S1A). These cultures are mixed cortical excitatory and inhibitory neurons as demonstrated by robust expression of the excitatory vesicular glutamate transporters marker (vGlut1), as well as the inhibitory glutamate decarboxylase 1 (GAD1) (Fig. S1B). We next assessed neuronal network activity by multielectrode array (MEA) analysis, which allows us to record electrical signals from thousands of neurons simultaneously (Fig. 1A). After 3 months in culture, both isogenic and P301L mutant neurons were spontaneously active and able to burst (Fig. 1B). Increases in neuronal firing were observed in mutant neurons at the single cell level, including measures such as firing rate and interspike interval (ISI) coefficient of variation (Fig. 1C). The network activity features were also altered by P301L tau. Mutant neurons fired longer bursts than the isogenic control (Fig. 1C). Although the network bursting rate did not change, the network bursting duration was enhanced in mutant neurons (Fig. 1B-C). These data indicate a hyperactive neuronal network in *MAPT* P301L neuronal cultures.

**Figure 1.**
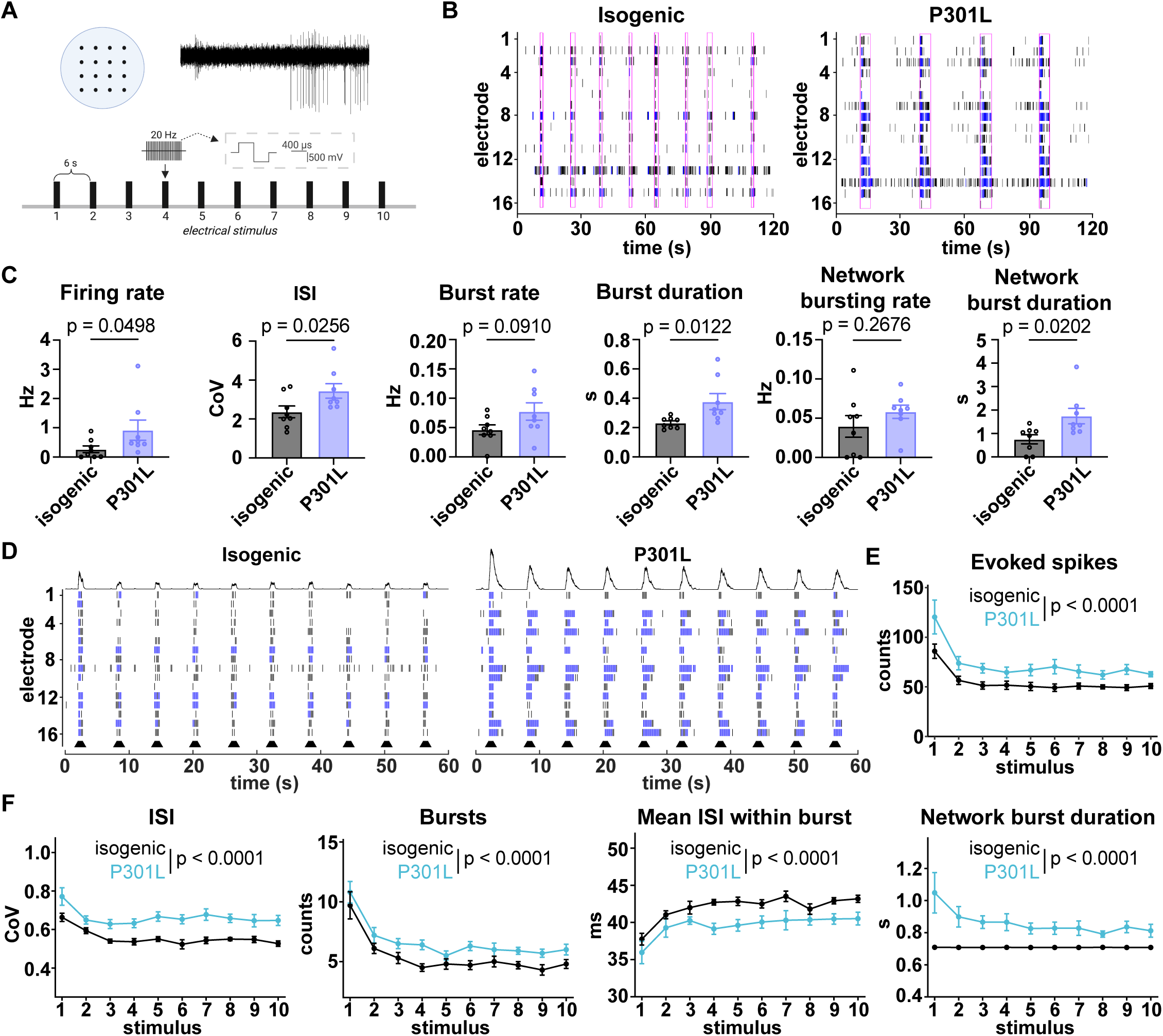
Neuronal and network hyperactivity in iPSC-iNs carrying *MAPT* P301L mutation. (A) An illustration of the multielectrode array (MEA) layout, a representative trace of spontaneous activity and the electrical stimulation protocol. (B) Representative raster plots of spontaneous activity recorded over 2 min in isogenic (left) and P301L (right) neuronal networks. (C) Quantification of spontaneous activity: firing rate, interspike interval (ISI), burst rate, burst duration, network bursting rate and network burst duration in isogenic control versus P301L neurons. Each data point represents the median of the well averages across one experimental repeat in which 12-24 wells per condition were included. Recordings were repeated 8 times. Unpaired t-test or Mann Whitney U test. (D) Representative raster plots of evoked activity recorded during repetitive electrical stimulation in isogenic (left) and P301L (right) neuronal networks. (E) Evoked spike counts across each successive stimuli in isogenic control and P301L networks. (F) Stimulus-evoked responses quantified as ISI, burst counts, mean ISI within burst, and network burst duration across ten consecutive stimuli in isogenic control and P301L cultures. Linear mixed-effects model followed by Benjamini-Hochberg post-hoc multiple comparisons. Data are mean +/-SEM; p values are indicated on each panel.

We also tested whether the network excitability was modulated by mutant tau and recorded evoked neuronal network activity. A brief high-frequency electrical stimulation can increase synaptic strength for prolonged periods [21,22]. To mimic this effect, we applied 10 trials of 20 Hz stimulation pulses within a 1-minute period. Both isogenic and mutant neurons showed adaptation in firing patterns after repetitive stimulations (Fig. 1D). During the stimulation, P301L neurons fired more spikes than the isogenic control (Fig. 1E). The evoked spike trains showed higher ISI coefficient variation in mutant neurons, suggesting a lower degree of spike regularity (Fig. 1F). In addition, mutant neurons fired more evoked spike bursts while the ISI within burst significantly decreased compared to the isogenic control (Fig. 1F). Finally, mutant neurons exhibited longer evoked network bursting duration compared to the isogenic control. These data indicate that P301L neurons exhibited higher excitability, although they remained plastic in response to repetitive stimulations. Collectively, P301L mutant tau increased neuronal and network burst activity in iPSC-derived cortical neurons.

### Neurophysiological measurements of network activity robustly predict tau P301L mutation

Next, we asked if neuronal and network activity changes measured by MEA could serve as a functional readout to differentiate neurons carrying *MAPT* P301L mutation from the isogenic control. We constructed a feature matrix that included quantitative measures of spontaneous activity at both cell and network levels. The cell-level features consisted of spike train statistics, such as firing rate, ISI coefficient of variation, synchrony index, and burst features such as burst frequency, burst duration, inter-burst interval (IBI), number of spikes per burst, and ISI within burst, among others. The network features included network burst frequency, network burst duration, ISI within network burst, ISI coefficient of variation within network burst, as well as inferred network burst features which were derived from binned spiking activity. Following construction of the feature matrix for each experimental repeat and normalization, we trained logistic regression models using either spike, burst, network burst or combined feature sets to classify mutant neurons and isogenic controls. The classifier was then used as an input to predict neuronal genotype (isogenic vs P301L). Performance of the classifiers was evaluated using repeated stratified two-fold cross-validation, in which half the dataset was used as training data and the other half was used as test data. The combined feature sets (spike + burst + network burst features) reached the highest accuracy for training the logistic regression model, compared to spike, burst or network burst feature sets alone (Fig. 2A). The combined feature classifier efficiently differentiated isogenic and P301L mutant neurons in the training dataset with an accuracy of 88.8%, and in the test dataset with an accuracy of 85.9% (Fig. 2B). The area under the receiver operating characteristic (ROC) curve reached 0.883, with sensitivity of 0.682 and specificity of 0.909 (Fig. 2C-D). The top weighted features in the trained model were ISI coefficient variation, burst duration, network burst duration interquartile range, ISI coefficient variation in network burst, number of spikes per network per channel, burst frequency, and related measures (Fig. 2E-F). Collectively, these data indicate that bursting and network-level activity features strongly predict the functional influence of *MAPT* P301L in human neurons.

**Figure 2.**
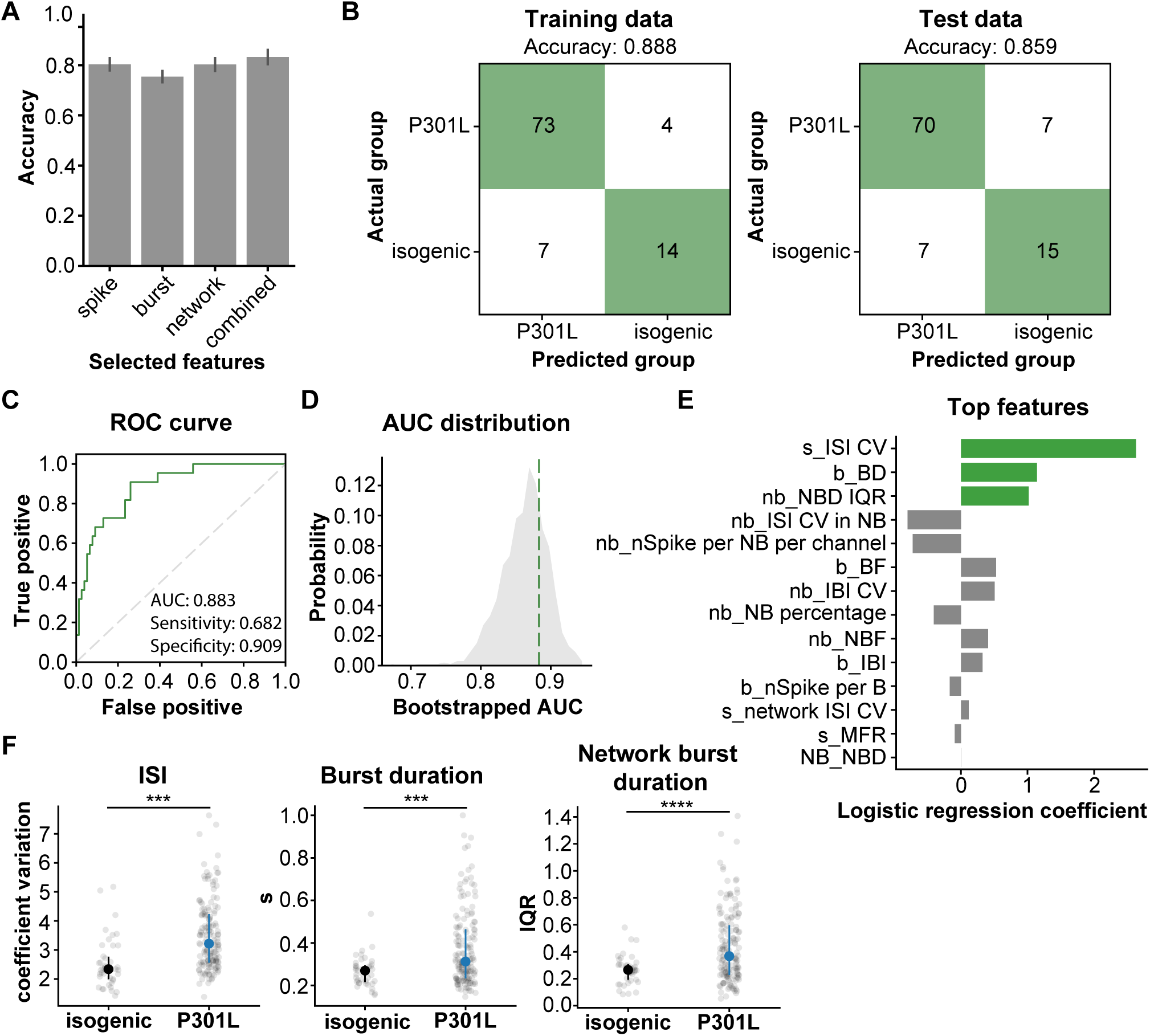
Electrophysiological signatures in multielectrode arrays robustly classify P301L network activity phenotypes. (A) Cross-validated classification accuracy of L1-regularized logistic regression models trained using spike, burst, network burst, or combined feature sets to classify P301L and isogenic control wells. Performance was evaluated using repeated stratified two-fold cross-validation. Bars show mean accuracy across folds; error bars indicate standard deviation. (B) Confusion matrices showing classification performance of the logistic regression model trained using the combined feature set following a stratified 50:50 train/test split. Left, predictions on the training dataset. Right, predictions on the held-out test dataset. Overall classification accuracy is indicated for each matrix. N = 43/154 wells for isogenic control/P301L mutants over 8 repeats. (C) Receiver operating characteristic (ROC) curve for genotype classification on the test dataset using combined features. The true positive rate is plotted against the false positive rate across classification thresholds, with the diagonal dashed line indicating chance-level performance. (D) Distribution of bootstrapped area under the ROC curve (AUC) values generated from repeated random train/test splits and model refitting. The vertical dashed line indicates observed AUC from the primary test set. (E) Logistic regression coefficients for the highest-weighted features from the L1-regularized model trained on the combined feature set. Positive coefficients indicate features associated with increased probability of P301L classification. ISI: inter-spike interval; BD: burst duration; NB: network bursting; NBD: network bursting duration; BF: bursting frequency; NBF: network bursting frequency, IBI: inter-burst interval; ISI: inter-spike interval; MFR: mean firing rate; s: spike features; b, burst features; nb: network burst features; CV: coefficient variation; IQR: interquartile range. (F) Representative features compared between genotypes. Individual wells are shown as gray dots. Black dots and line showed the summary statistics of the median and interquartile range (25% - 75%). N = 43/154 wells for isogenic control / P301L mutants over 8 repeats. Median test, p-values = 1.7e-4/1.7e-4/8.7e-6 for ISI CV/burst duration/network burst duration IQR, respectively.

### Differential impact of pathological tau on the firing properties of excitatory vs. inhibitory neurons

Bursting activity within a network is critically governed by a complex interplay of intrinsic cellular mechanisms and network dynamics. The balance between excitation and inhibition uniquely shapes burst characteristics [23]. We speculated that excitation and/or inhibition might change in neurons with mutant tau. To test this hypothesis, we sorted the spontaneous spikes according to their waveforms and grouped the single-unit spikes based on cross-correlograms of the spike timing of single neuronal activity. The activity of excitatory units represented putative excitatory neurons (Fig. 3A) while the activity of inhibitory units represented putative inhibitory neurons (Fig. 3F). In mutant neurons, the firing rate of excitatory units increased (Fig. 3B). More variation in spike train dynamics, as indicated by increased ISI coefficient of variation, was observed in the presence of mutant tau (Fig. 3C). In addition, the variation of burst duration also increased in excitatory units (Fig. 3D). Consistently, the interburst interval (IBI) decreased, compared to the isogenic control (Fig. 3E). Fewer changes were observed in inhibitory units in the presence of mutant tau. The firing rate was similar in isogenic vs mutant neurons (Fig. 3G), whereas ISI variation increased (Fig. 3H), and the burst of inhibitory units decreased in mutant neurons (Fig. 3I). These data indicate that the functional influence of pathological tau varies across different neuronal subtypes.

**Figure 3.**
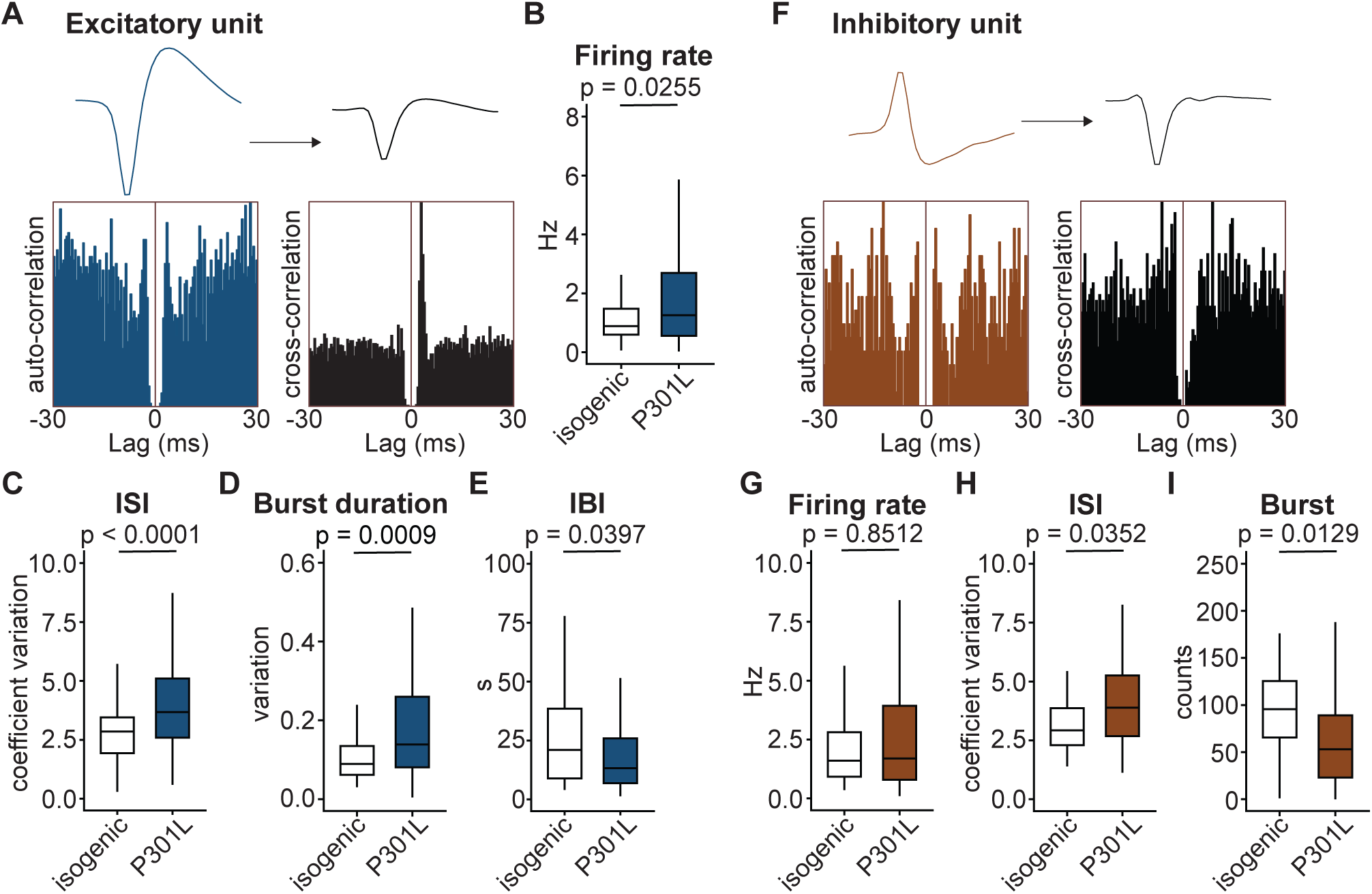
Differential changes in firing patterns of putative excitatory vs. inhibitory neurons in P301L mutant neurons. Single-unit spike waveforms of the spontaneous activity were sorted and grouped according to their shape and spike-timing on the cross-correlation graph. (A) Representative graphs of a group of putative excitatory units. Left are an exemplar waveform (top) and auto-correlation graph (bottom) of the excitatory units. Right shows a group of units that are likely activated by the excitatory units with different waveform (top) and cross-correlation graph (bottom). Quantification of the (B) firing rate, (C) ISI coefficient of variation, (D) burst duration, (E) IBI, and (F) IBI coefficient of variation of putative excitatory units from all 8 recordings. (G) Representative graphs of an isolated group of putative inhibitory units. Left are an exemplar waveform (top) and auto-correlation graph (bottom) of the inhibitory units. Right shows a group of units that likely are suppressed by the inhibitory units with different waveform (top) and cross-correlation graph (bottom). Quantification of the (H) firing rate, (I) ISI coefficient of variation, and (J) burst counts of putative inhibitory units from all 8 recordings. IBI: inter-burst interval; ISI: inter-spike interval. Unpaired t-test or Mann Whitney U test; p values are indicated on each panel.

To explore the differential functional influence of tau pathology in each neuronal subtype, we genetically labeled excitatory neurons or inhibitory neurons, followed by patch-clamp experiments to assess their electrophysiological properties (Fig. 4A and 4D). First, we examined the intrinsic properties of action potential (AP) firing. Excitatory neurons with mutant tau had similar passive membrane electrophysiological properties including resting membrane potential, input resistance and membrane capacitance as excitatory isogenic neurons (Table 2). By injecting hyperpolarizing and depolarizing currents, mutant excitatory neurons fired APs at a higher frequency than the isogenic control (Fig. 4B-C). Similar analyses were conducted in inhibitory neurons. As in excitatory neurons, mutant tau did not change the resting membrane potential, input resistance, and membrane capacitance in inhibitory neurons (Table 2). However, in contrast to the excitatory neurons, mutant inhibitory neurons fired APs at a lower frequency than the isogenic control (Fig 4E-F). We verified these findings in neurons differentiated from a second iPSC line with *MAPT* P301L mutation and its isogenic control and observed similar results (Fig. S1C). Taken together, these results indicate that *MAPT* P301L differentially affects intrinsic firing modes of excitatory neurons and inhibitory neurons, further indicating distinct mechanisms underlying the pathological influence of *MAPT* P301L in excitatory vs. inhibitory neurons.

**Figure 4.**
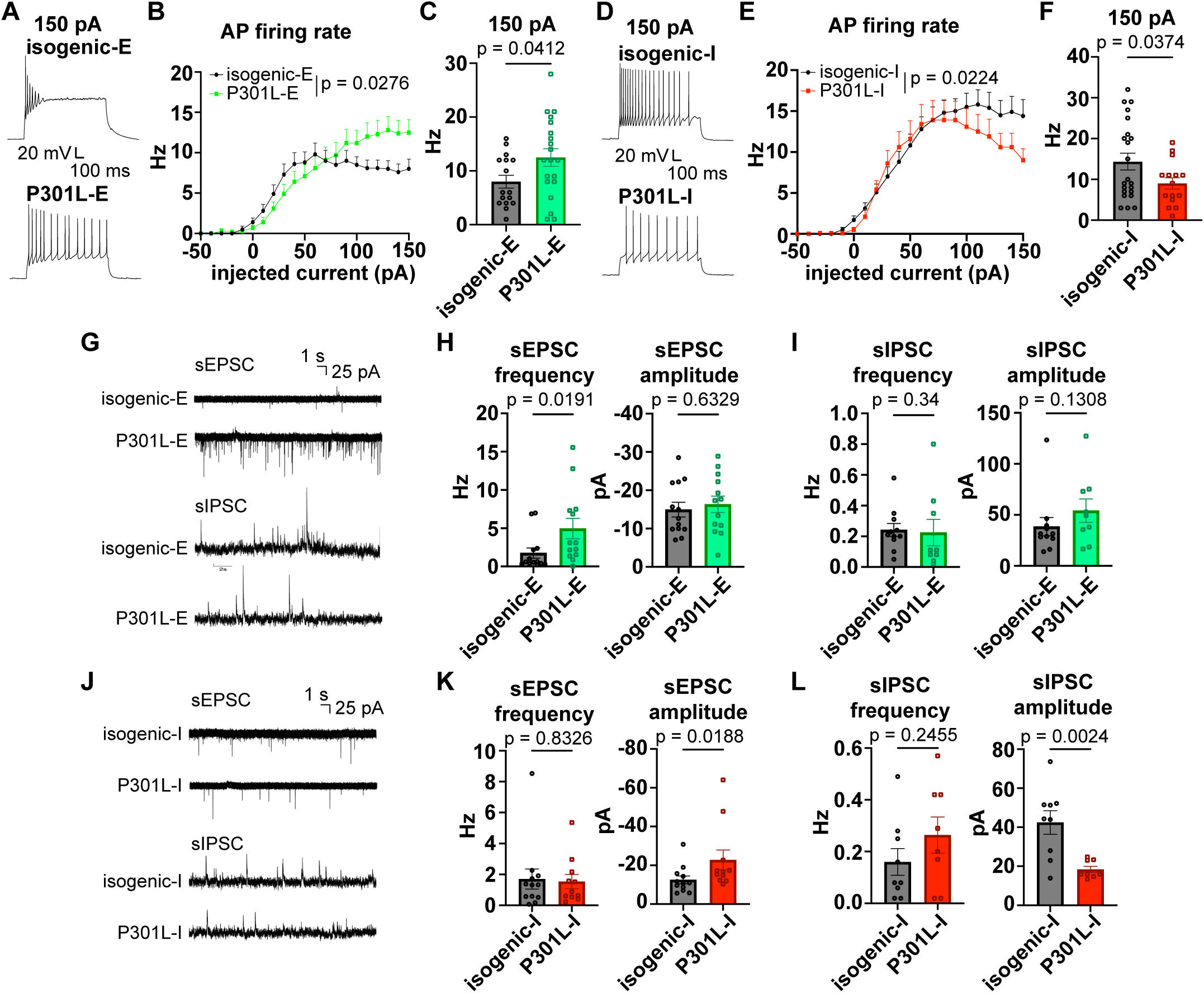
Mutant P301L differentially modulates neuronal activity, enhancing excitatory and suppressing inhibitory neuron activities. (A) Representative whole-cell current-clamp traces of action potentials evoked by 150 pA current injection in isogenic control (isogenic-E) and P301L (P301L-E) excitatory neurons. (B) Action potential (AP) firing rate as a function of injected current (F/I curve) in isogenic-E and P301L-E neurons. Two-way ANOVA. (C) Firing rate at 150 pA current injection in excitatory neurons. Unpaired t-test. (D) Representative whole-cell current-clamp traces of action potentials evoked by 150 pA current injection in isogenic control (isogenic-I) and P301L (P301L-I) Inhibitory neurons. (E) F/I curves in isogenic-I and P301L-I neurons. Two-way ANOVA. (F) Firing rate at 150 pA current injection in inhibitory neurons. Unpaired t test. (G) Representative spontaneous EPSC (sEPSC) and IPSC (sIPSC) recordings from excitatory neurons. (H) The frequency but not amplitude of sEPSC significantly increased in P301L-E neurons compared to isogenic-E neurons. (I) Neither frequency nor amplitude of sIPSC differed between P301L-E vs control-E. (J) Representative spontaneous EPSC (sEPSC) and IPSC (sIPSC) recordings from inhibitory neurons. (K) The frequency of sEPSC was comparable between P301L-I and isogenic-I neurons. However, the amplitude of sEPSC significantly increased in P301L-I neurons. (L) The frequency did not differ between P301L-I and isogenic-I neurons, whereas its amplitude significantly decreased in P301L-I compared to control-I. Data are mean +/-SEM; p-values by unpaired t-test or Mann-Whitney U test. A total of 8-13 cells from at least two coverslips were included.

We also assessed synaptic transmissions by analyzing post-synaptic currents by voltage clamp (Fig. 4G and 4J). In excitatory neurons, the frequency but not amplitude of spontaneous excitatory post-synaptic current (sEPSC) increased in mutant neurons compared to the isogenic control (Fig. 4H). The same pattern was observed for miniature excitatory post-synaptic current (mEPSC), suggesting enhanced presynaptic transmission (Fig. S2A). In contrast, neither frequency nor amplitude changed in inhibitory post-synaptic current (sIPSC and mIPSC) in excitatory neurons (Fig. 4I and Fig. S2B). In inhibitory neurons, the frequency of sEPSC was comparable between mutant and isogenic control (Fig. 4K). However, the amplitude of sEPSC increased in mutant inhibitory neurons (Fig. 4K). The frequency of mEPSC remained similar between mutant and isogenic neurons, while the amplitude of mEPSC decreased in mutant inhibitory neurons (Fig. S2C). In addition, the frequency of sIPSC did not change, whereas the amplitude of sIPSC decreased (Fig. 4L). The same trend was observed in mIPSC but it did not reach statistical significance (Fig. S2D). These data altogether indicate altered synaptic transmission in mutant neurons. P301L tau increased excitation in excitatory neurons, whereas both excitation and inhibition changed in inhibitory neurons.

### Altered excitatory vs inhibitory ratio and tau level in neurons with pathological tau

We then compared the expression of inhibitory vs excitatory neuronal markers between the isogenic control and mutant neurons. Decreased mRNA expression of inhibitory markers GAD1 and vGAT was observed in mutant neurons compared to the isogenic controls (Fig. S3A). In contrast, the excitatory markers vGlut1 and vGlut2 were upregulated in mutant neurons at the transcriptional level (Fig. S3B). Similar changes were observed at the protein level by western blotting. GAD1 and the pan neuronal marker NeuN decreased, while CaMK2a, a general glutamatergic neuronal marker, increased in tau mutant neurons, compared to the isogenic control (Fig. 5A). Furthermore, a significant reduction of GABA positive neurons was observed in mutant neurons by immunostaining of GABA and MAP2 (Fig. 5B). However, no change in the proportion of vGlut1 positive neurons was observed (Fig. 5C). These results indicate altered cellular composition in neurons carrying *MAPT* P301L mutation.

**Figure 5.**
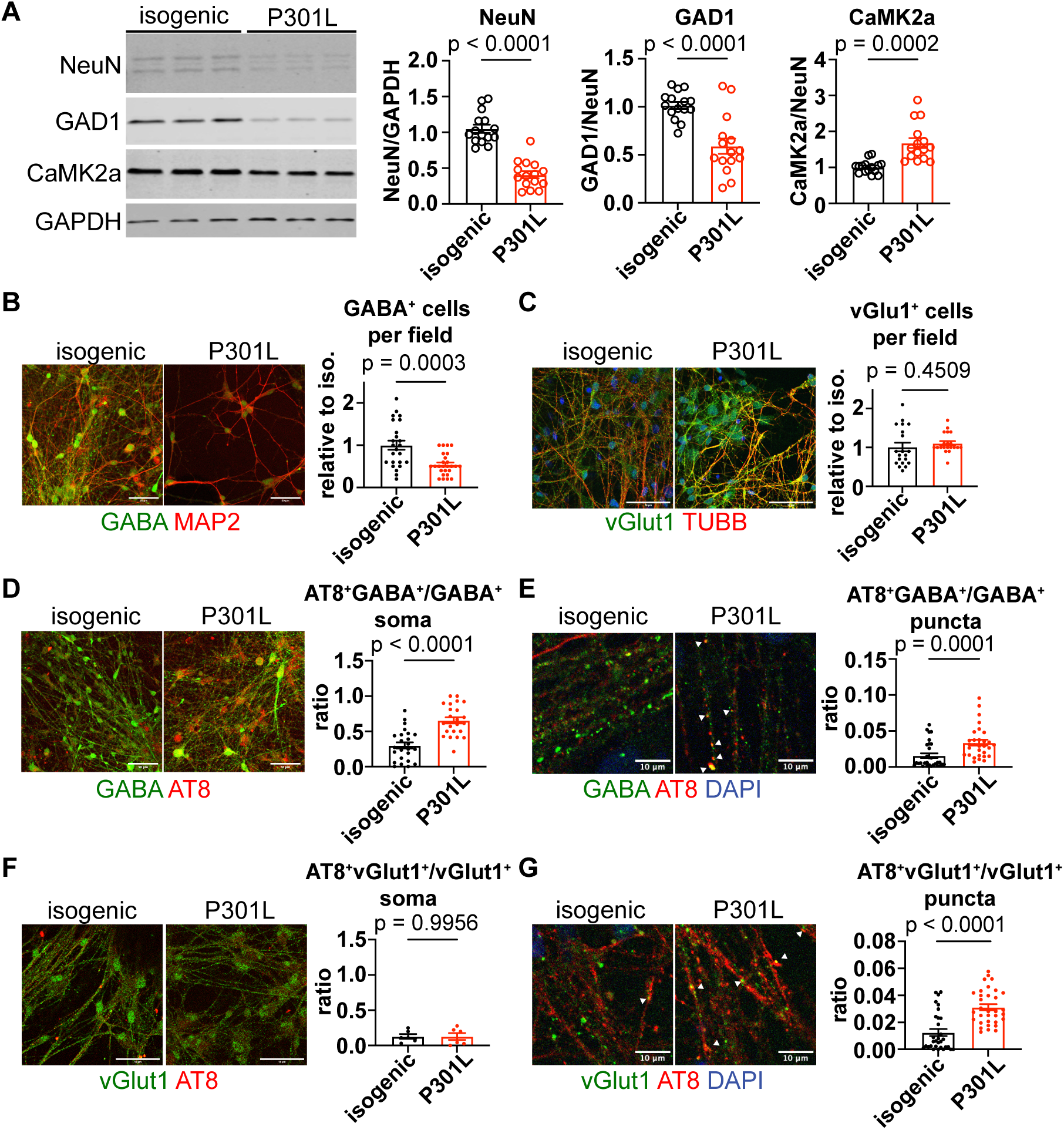
Differential changes in excitatory vs. inhibitory neuronal markers and selective accumulation of p-tau in inhibitory neurons with mutant P301L tau. (A) Representative Western blots of inhibitory marker GAD1, excitatory marker CaMK2a and pan-neuronal marker NeuN. GAPDH is a loading control. Quantifications shown right of the blots were from four independent batches of iPSC-iN differentiation. Relative expression was normalized to NeuN or GAPDH, and compared to the isogenic control. Unpaired t test, p value as indicated. (B) Representative immunostaining of GABA^+^ cells (GAD1/MAP2, left) and vGluT1^+^ cells (vGluT1/TUBB3, right). GABA and MAP2 positive or vGlut1 and TUBB positive cells per imaging field were quantified. A total of 20-26 fields were counted for each line and normalized to the isogenic control. (C) Representative images and quantification of AT8^+^ (p-tau) signal in GABA^+^ neurons, expressed as the ratio of AT8^+^GABA^+^ / GABA^+^ soma or AT8^+^GABA^+^ / GABA^+^ puncta. (D) Representative images and quantification of AT8^+^ signal in vGluT1^+^ neurons, expressed as the ratio of AT8^+^vGluT1^+^ / vGluT1^+^ soma or AT8^+^vGluT1^+^ / vGluT1^+^ puncta. Data are mean +/-SEM with p values shown. Unpaired t test. B-D and F were taken under a 20x objective, scale bar, 50 µm; E and G were taken under a 63x objective, scale bar, 10 µm.

We next asked if tau expression is different in each neuronal subtype. The major tau isoform expressed in these neurons is 3R tau; however, the level of 4R tau gradually increased over the course of differentiation (Fig. S4A). Western blot analysis revealed the presence of both 3R and 4R tau at day 120 post differentiation, and 3R tau was the predominant isoform (Fig. S4B-C). Next, we assessed the levels of phosphorylated tau (ptau). All ptau and total tau increased in mutant neurons compared to isogenic controls (Fig. S4D). When examined in subpopulations by co-immunostaining, tau pS202/T205 accumulated in the soma of GABA positive inhibitory neurons, but not in excitatory neurons (Fig. 5D and 5F). An increase in ptau containing inhibitory soma, but not excitatory soma was observed in mutant P301L neurons compared to their isogenic control (Fig. 5D and 5F). Notably, punctate localization of ptau along both inhibitory and excitatory neuronal processes was observed (Fig. 5E and 5G), suggesting synaptic localization of ptau. The ptau positive GABA positive or vGlut1 positive puncta increased in mutant neurons compared to the isogenic control (Fig. 5E and 5G). These data indicate phosphorylated tau selectively accumulates in inhibitory neurons, as well excitatory and inhibitory synapses.

### Contribution of excitatory synaptic transmission in enhanced neuronal network excitability with pathological tau

Given increased vGlut1/2 expression, enhanced excitatory synaptic transmission and synaptic localization of ptau in excitatory neurons with tau P301L mutation, we first tested if increased excitatory synaptic transmission could be the driving force of neuronal network hyperexcitability. The frequency of both spontaneous and miniature EPSCs increased in mutant excitatory neurons (Fig. 4H and S2A), indicating a presynaptic upregulation of glutamate release. By applying the AMPA receptor antagonist NBQX to mutant neurons, both the firing rate and ISI coefficient of variation of mutant neurons decreased (Fig. S5A). During high frequency stimulation, the evoked spike counts decreased, and network burst duration reverted to the isogenic level by NBQX treatment (Fig. S5B). Similar treatments were also performed by applying the NMDA receptor antagonist APV. In contrast to NBQX treatment, no change in firing rate or ISI coefficient variation was observed in mutant neurons (Fig. S5C). In addition, APV treatment did not alter the evoked spike counts or network burst duration of mutant neurons (Fig. S5D). Taken together, these data indicate that AMPA receptor mediated excitatory synaptic transmission contributes to enhanced network excitability caused by P301L mutant tau.

### Pathological tau differentially modulates tau interactomes in excitatory vs. inhibitory neurons

To further dissect the mechanisms underlying the differential influence of mutant tau in excitatory and inhibitory neurons, we selectively expressed Flag-tagged full length 2N4R wild-type (TauWT) or 2N4R tau carrying the P301L mutation (TauP301L) in isogenic neurons (Fig. 6A). TauWT or TauP301L were exclusively expressed in either excitatory or inhibitory neurons (Fig. 6B). Following Flag immunoprecipitation, TauWT or TauP301L were enriched and subjected to mass spectrometry to identify steady-state tau interacting proteins. First, we analyzed proteins enriched in TauWT interactomes from excitatory vs inhibitory neurons (Table 3). Of the putative tau interacting proteins, 134 were enriched in both excitatory and inhibitory neurons (Fig. 6C). Gene ontology (GO) enrichment analyses revealed that the molecular function of these proteins, largely involves cytoskeleton structural networks, as well as protein folding, cell adhesion molecule binding and ubiquitin protein ligase binding (Fig. 6D) as previously reported [24]. There were 124 proteins uniquely enriched in the TauWT interactome of excitatory neurons (Fig. 6C). GO analysis revealed that these proteins are primarily involved in ribonucleoprotein complex assembly, RNA splicing, RNA export from the nucleus and protein deacetylation (Fig. 6E). Cellular localizations of these proteins include spliceosomes, extracellular exosome, ribonucleoprotein complex, nucleosome remodeling and deacetylase complex, and nuclear speck (Fig. 6F), consistent with previous reports on the nuclear localization of tau [25]. There were 58 proteins uniquely enriched in the TauWT interactome of inhibitory neurons (Fig. 6C). GO analysis demonstrated that these proteins are primarily involved in cation channel activity, such as Na^+^, K^+^ and Ca^2+^ transporters, and protein kinase inhibitor activity (Fig. 6G). Cellular localizations of these proteins include dendritic spine neck, postsynaptic and dendritic cytoskeleton, and extracellular exosome (Fig. 6H).

**Figure 6.**
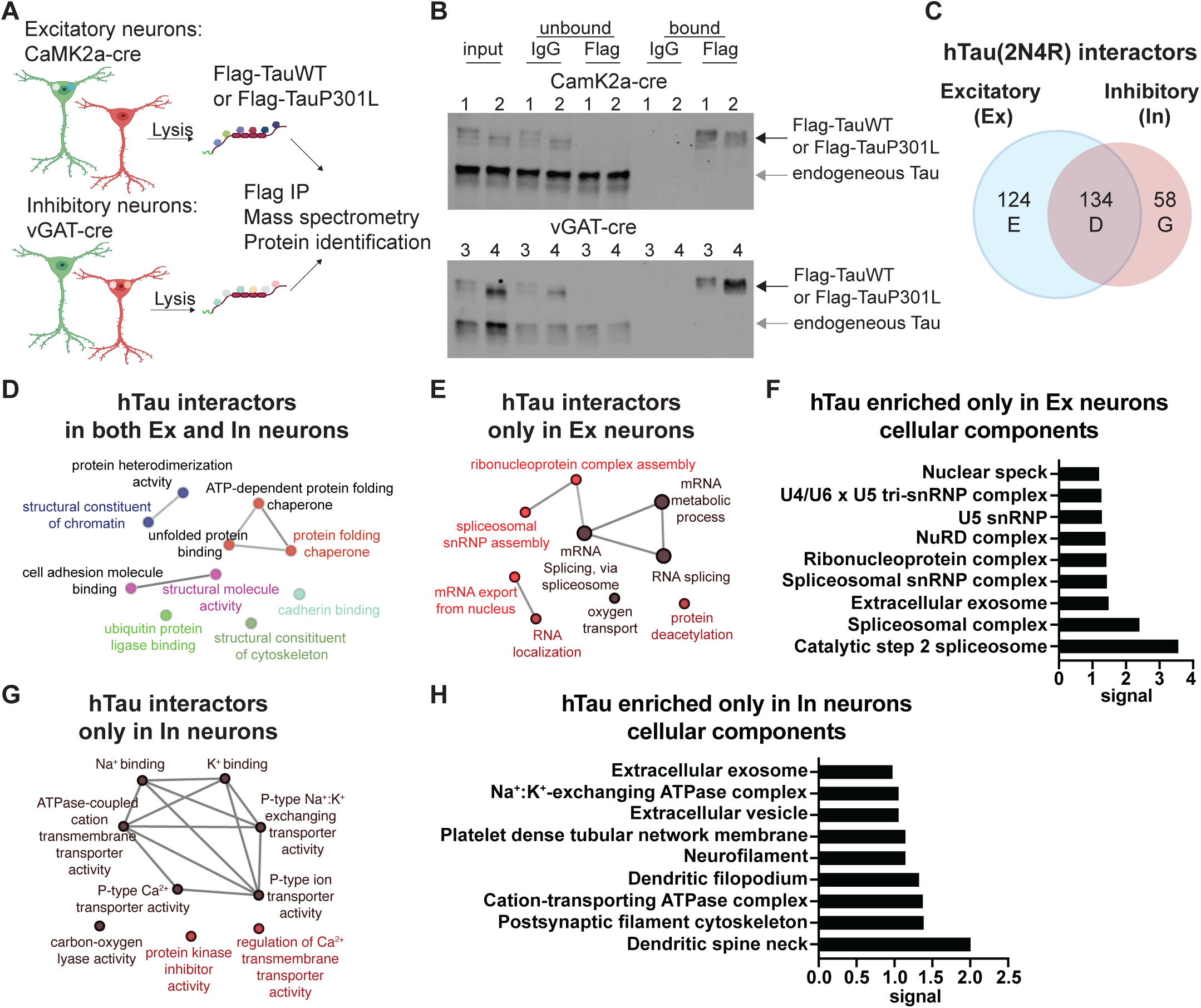
Cell-type specific tau interactomes in excitatory vs. inhibitory neurons. (A) Graph illustration of the workflow for protein identification after Flag affinity purification. Flag-TauWT or Flag-TauP301L were expressed in either excitatory or inhibitory neurons under cre-dependent conditions in the isogenic background. (B) Western blots of Flag-TauWT or Flag-TauP301L by HT7 tau antibody. Black arrows denote exogenous tau; grey arrows indicate the endogenous tau. Lanes 1 and 3 are Flag-TauWT expression; lanes 2 and 4 are Flag-TauP301L expression. (C) Venn diagram of proteins identified as common interactors in both excitatory (Ex) and inhibitory (In) neurons, or unique interactors in Ex or In neurons. (D) ClueGO molecular function pathway enrichment of the 134 proteins that are common tau interactors in both Ex and In neurons. (E) ClueGO molecular function pathway enrichment of the 124 proteins that are unique tau interactors in Ex neurons. (F) STRING analysis of the enriched cellular components of the unique tau interactors in Ex neurons. (G) ClueGO molecular function pathway enrichment of the 58 proteins that are unique tau interactors in In neurons. (H) STRING analysis of enriched cellular components of the unique tau interactors in In neurons.

To identify tau interactomes altered by tau P301L mutation in each neuronal subtype, we compared TauWT interacting proteins with TauP301L interacting proteins and focused on interactions that were suppressed or enhanced by the P301L tau. In excitatory neurons, TauP301L had decreased interaction with 14 proteins compared to TauWT (Fig. 7A). Only 7 proteins showed enhanced interaction with TauP301L (Fig. 7A). These proteins are involved in glutamate signaling, actin and microtubule binding, RNA splicing and intracellular organelle trafficking (Table 4). In inhibitory neurons, TauP301L had enhanced interaction with 20 proteins and led to decreased interaction with 9 proteins compared to TauWT (Fig. 7B). These proteins include cytoskeletal scaffolding proteins, molecular chaperones, endoplasmic reticulum proteins, mitochondrial protein, and core and regulatory subunits of the 26S proteasomes (Table 4).

**Figure 7.**
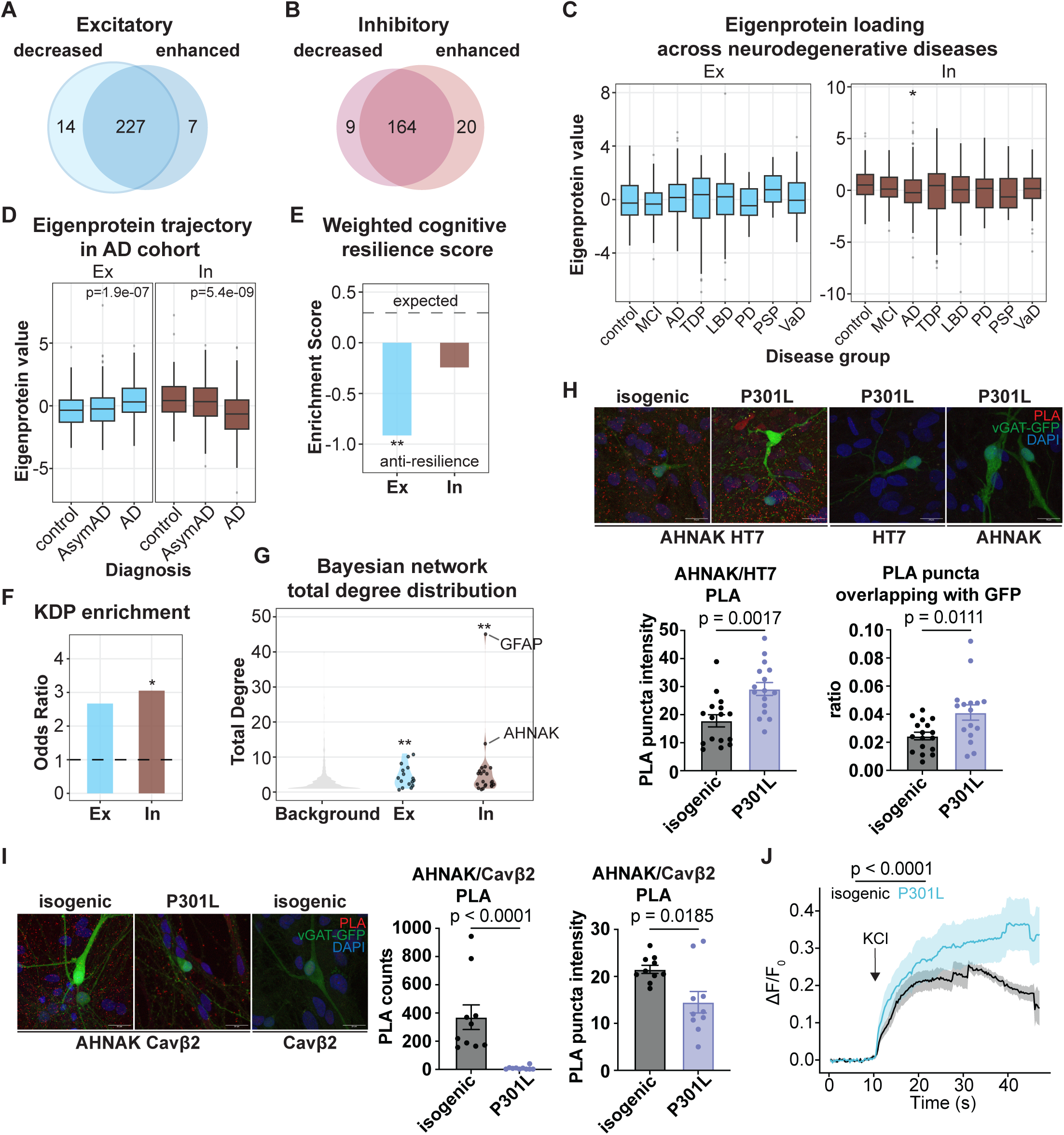
**Altered tau interactors by P301L tau in excitatory or inhibitory neurons associate with AD progression. (**A) Vinn diagram of tau interactors with decreased or increased interaction with P301L tau in excitatory neurons. (B) Vinn diagram of tau interactors with decreased or increased interaction with P301L tau in inhibitory neurons. (C) Eigenprotein trajectory of the tau interacting proteins altered by P301L tau in excitatory neurons (Ex) or inhibitory neurons (In) across six neurodegenerative diseases from Shrestha et al 2026^30^. Wilcoxon rank-sum test with Benjamini-Hochberg post hoc multiple comparisons. (D) Eigenprotein trajectory of the tau interacting proteins altered by P301L tau in Ex or In across AD stages using ROSMAP and Banner cohorts. Kruskal-Wallis test with post-hoc Benjamini-Hochberg correction. (E) Weighted enrichment score for the tau interacting proteins altered by P301L tau in Ex or In in the anti-cognitive resilience protein network from the ROSMAP cohort. Fisher’s exact test and a weighted permutation enrichment. (F) Enrichment of the tau interacting proteins altered by P301L tau in Ex or In in key driver protein (KDP) network that is associated with AD progression identified by protein co-expression and Bayesian causal network analyses in Wang et al 2025^28^. Fisher’s exact test. (G) Bayesian network analysis of the total degree distribution for the tau interacting proteins altered by P301L tau in Ex or In. Wilcoxon rank-sum tests. (H) Proximity ligation assay (PLA) validation of AHNAK and tau (HT7) interaction in inhibitory neurons. Representative immunostainings in isogenic control and P301L neurons are shown above. Scale bar: 50 μm. Quantifications of PLA puncta and the overlapping between PLA and inhibitory markers are shown below. Unpaired t test. (I) PLA of AHNAK and CACNB2 interaction in isogenic and P301L inhibitory neurons. Representative immunostainings are shown at left, and quantifications of PLA signals are shown at right. Scale bar: 50 μm. Unpaired t test. (J) Ca^2+^ influx upon 30 mM KCl depolarization in isogenic and P301L neurons. Linear mixed-effects model, a group-by-time interaction was used to assess whether the temporal response differed between groups. Data are mean +/-SEM; p values are shown on each panel.

Next, we asked if tau interacting proteins altered by P301L overlap with AD or FTD-associated protein expression modules (excitatory list: 20 proteins; inhibitory list: 29 proteins). We leveraged four independent human brain proteomic resources across neurodegenerative diseases [26–30]. Shrestha et al 2026 reported multilayer proteomic signatures from whole proteome, detergent-insoluble proteome and post-translational modifications across multi-disease cohorts including AD, FTD with TDP proteinopathy (FTD-TDP), progressive supranuclear palsy with tauopathy (PSP/tauopathy), Lewi body dementia (LBD), Parkinson’s disease (PD), vascular dementia (VaD), and healthy controls. Johnson et al 2022 analyzed protein co-expression network using the Religious Orders Study and Memory and Aging Project (ROSMAP) and the Banner Sun Health Research Institute (Banner) cohorts of full AD continuum. To map our tau interacting protein lists that altered by P301L tau, we computed a list-level eigenprotein against each dataset, and sign-aligned between these two datasets so that positive values denote higher aggregate expression of the list. In Shrestha 2026 dataset, our tau interactors altered in excitatory neurons showed a trend of elevation across the tauopathy and synucleinopathy spectrum, with the largest trend of separation in PSP/tauopathy (control vs. PSP/tauopathy, p = 0.082); tau interactors altered in inhibitory neurons decreased most strongly in AD (control vs. AD, p = 0.0020, FDR = 0.014) and trending downward in PSP/tauopathy (Fig. 7C). Thus, tau interactors altered by P301L in excitatory neurons and inhibitory neurons showed opposing eigenprotein trajectories in the Shrestha 2026 cohort. This cross-disease pattern was recapitulated in the Johnson 2022 ROSMAP and the Banner cohorts, where the eigenprotein of both lists showed significant diagnosis-dependent variation (Fig. 7D). Across the AD pathological continuum: tau interactors altered in excitatory neurons rose from control through asymptomatic AD (AsymAD) to AD, while tau interactors altered in inhibitory neurons fell from control through AsymAD to AD, with changes in both lists reaching significance in AD vs control (excitatory list, p = 4.5×10^−6^; inhibitory list, p = 1.0×10^−7^) (Fig. 7D). This shift indicates that the tau interactors in our study play a role in tracking initiation and development of AD pathology.

Wingo et al 2019 performed a proteome-wide association study in the Banner and Baltimore Longitudinal Study of Aging (BLSA) cohorts, correlating protein abundance with the longitudinal rate of cognitive decline [26]. Using a weighted enrichment score that aggregates per-protein effects on cognitive resilience, we mapped the tau interactors altered by P301L in excitatory neurons or inhibitory neurons to this dataset. The weighted score of the excitatory list was significantly biased toward anti-resilience network (weighted score −0.914, Z = −2.42, p = 0.0075), driven by four anti-resilience members (ANXA1, CSTB, HIST1H4A, UGP2; odds ratio = 3.89, p = 0.035); the weighted score of the P301L altered inhibitory list showed the same anti-resilience directionality but did not reach significance, with GFAP, AHNAK, and HSPB1 contributing anti-resilience effects and ACTN4 as a pro-resilience member (Fig. 7E). Finally, we compared our lists to Wang et al 2025 which combined protein co-expression and Bayesian causal network approaches to identify key protein subnetwork and drivers of AD progression. Notably, 5 tau interactors altered by P301L in inhibitory neurons were enriched among key driver proteins (KDPs) - ACTG1, GFAP, AHNAK, RPL12, TUBB3 (odds ratio = 3.05, p = 0.038), whereas tau interactors altered by P301L in excitatory neurons were not (Fig. 7F). Total-degree analysis of the Bayesian network further distinguished the two lists: tau interactors altered in inhibitory neurons were highly connected, led by GFAP and AHNAK, whereas tau interactors altered in excitatory neurons were modestly connected (Fig. 7G). Taken together, tau interactors altered by P301L in excitatory neurons represent an anti-resilient program enriched in RNA-splicing and cytoskeletal processes whose increase tracks with AD progression. In contrast, tau interactors altered by P301L in inhibitory neurons change in an opposite direction whose decline tracks with AD progression, with members as causal hubs identified in the AD network by human studies.

### Pathological tau interacts with AHNAK and modulates calcium influx in inhibitory neurons

Among the list of tau interactors altered by P301L tau, neuroblast differentiation-associated protein AHNAK is one of the key driver proteins of the AD network identified by Wang et al 2025. AHNAK has been shown to directly regulate Ca^2+^ influx through the regulatory β subunit of L-type voltage-gated Ca^2+^ channel (VGCC) [31]. We next validated this candidate tau interactor and assessed if tau-AHNAK interaction was involved in regulating neuronal activities. The interaction between AHNAK and tau was confirmed by proximity ligation assay (PLA) in both P301L mutant and isogenic neurons (Fig. 7H). Negative controls including only one antibody, confirmed the specificity of PLA signals. The PLA signal increased in P301L mutant neurons compared to the isogenic control (Fig. 7H). Additionally, the ratio of PLA puncta in inhibitory neurons was higher in mutant neurons than in isogenic neurons (Fig. 7H). These results provide direct evidence that tau interacts with AHNAK, and their interaction is enhanced by tau P301L mutation. AHNAK slows cardiac VGCC inactivation through a direct interaction with the regulatory Cavβ2 subunit and leads to increased Ca^2+^ current influx upon depolarization [32,33]. In P301L mutant inhibitory neurons, the interaction between AHNAK and neuronal Cavβ2 decreased in the presence of P301L tau (Fig. 7I). Given that AHNAK acts as a physiological brake on Ca^2+^ influx [31], we hypothesized that enhanced TauP301L-AHNAK interaction and decreased AHNAK-Cavβ2 leads to increased Ca^2+^ in mutant neurons. To test this hypothesis, we performed Ca^2+^ imaging on inhibitory neurons expressing the genetically encoded calcium indicator GCaMP7s. Upon 30 mM KCl stimulation which depolarized the membrane potential and activated VGCCs, an increase in intracellular Ca^2+^ was observed in both isogenic and mutant inhibitory neurons (Fig. 7J). However, mutant inhibitory neurons had much more Ca^2+^ influx than the isogenic control (Fig. 7J). This data is consistent with our prediction and indicate that enhanced P301L tau and AHNAK interaction may associate with enhanced Ca^2+^ influx during neuronal depolarization in inhibitory neurons.

### Enhanced Ca^2+^ influx through voltage-gated calcium channel activity in inhibitory neurons contributes to enhanced neuronal network activity in P301L tau mutant neurons

Next, we tested if enhanced Ca^2+^ in mutant neurons altered neuronal excitability and network activity. First, we performed whole-cell patch clamp experiments to assess if excess intracellular Ca^2+^ alters neuronal excitability. Cell-impermeable BAPTA was included in the internal solution to lower free intracellular Ca^2+^. In the presence of BAPTA, mutant P301L excitatory neurons fired less evoked action potentials (Fig. 8A). In contrast, mutant P301L inhibitory neurons exhibited enhanced excitability in the presence of BAPTA (Fig. 8B). Taken together, excess intracellular Ca^2+^ indeed altered neuronal excitability in excitatory vs inhibitory neurons differently, which may lead to enhanced neuronal network activity in mutant neurons.

**Figure 8.**
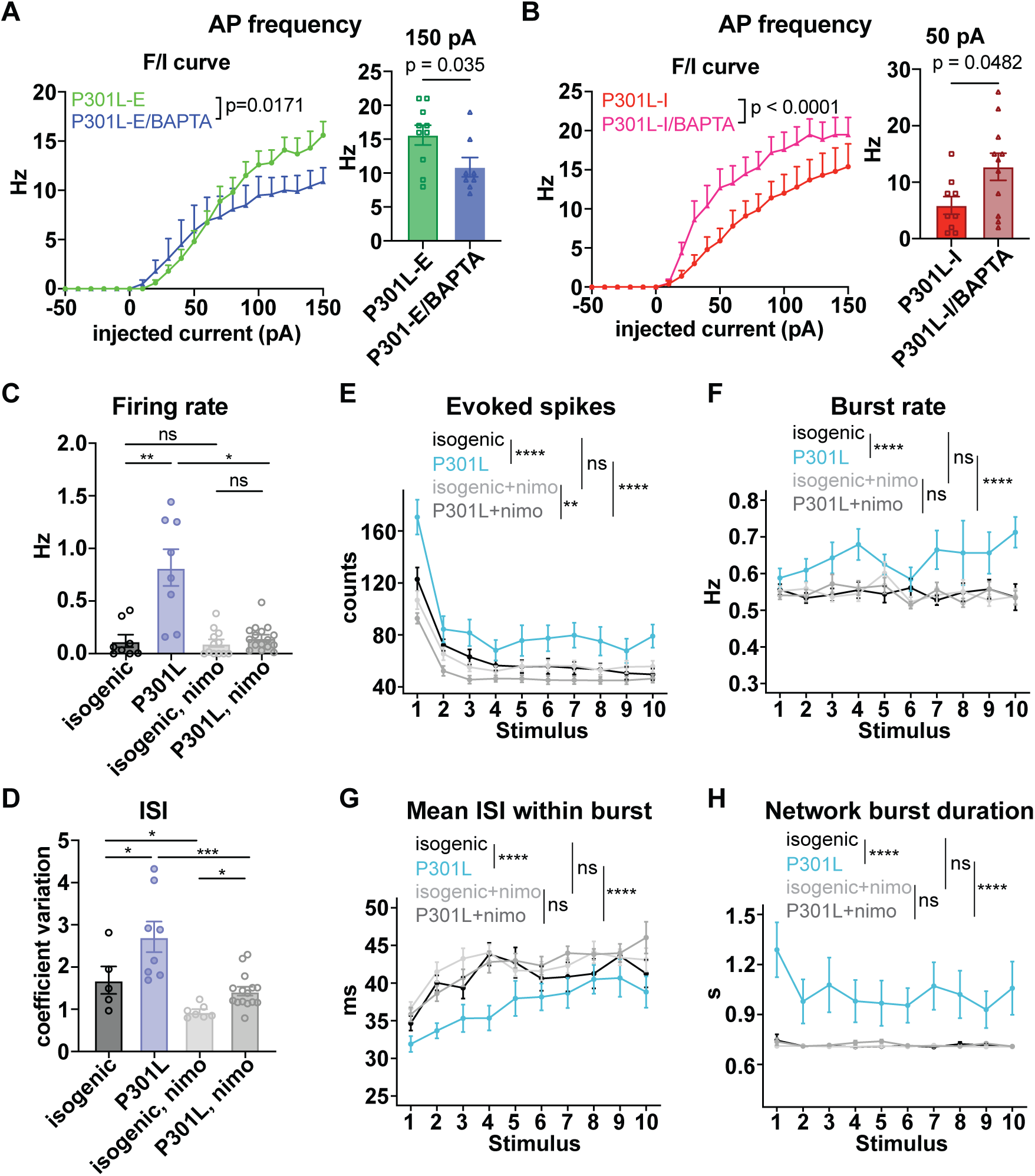
Elevated Ca^2+^ through voltage gated Ca^2+^ channel (VGCC) underlies increased excitability and network hyperactivity of P301L neurons. (A) F/I curves in excitatory P301L neurons with and without BAPTA in the internal solution. AP frequency at the 150 pA current injection is shown on the right. (B) F/I curves in inhibitory P301L neurons with and without BAPTA in the internal solution. AP frequency at the 50 pA current injection is shown on the right. (C-D) MEA recordings of (C) spontaneous firing rate and (D) ISI coefficient of variation of isogenic control and P301L neurons in the absence or presence of VGCC blocker nimodipine. (E-H) MEA recordings of (E) evoked spikes, (F) burst rate, (G) mean ISI within burst, and (H) network burst duration during electrical stimulation. (A-B) Two-way ANONA followed by Sidak’s multiple comparisons test on the left and unpaired t test on the right, (C-D) One-way ANOVA followed by Tukey’s post-hoc test. (E-F) and (G - H) Linear mixed-effects model followed by Tukey’s post-hoc test. Data are mean +/-SEM; ns, not significant; *p < 0.05, **p < 0.01, ***p < 0.001, ****p < 0.0001.

To further investigate if L-type VGCC played a role in mediating enhanced Ca^2+^ influx to cause neuronal network hyperactivity, we applied nimodipine to block L-type VGCC function during MEA recording. Nimodipine significantly decreased the spontaneous firing rate in mutant P301L neurons, however, it did not affect the firing rate of isogenic neurons (Fig. 8C). The ISI coefficient variation significantly decreased in both isogenic control and mutant neurons in the presence of nimodipine, compared to their corresponding untreated controls (Fig. 8D). During high frequency electrical stimulation, the evoked spike counts in mutant neurons were reduced by nimodipine treatment, however, the isogenic control was not affected (Fig. 8E). Additionally, nimodipine corrected the enhanced evoked burst rate in mutant neurons without any effect on isogenic neurons (Fig. 8F). Moreover, mean ISI within burst was increased in mutant neurons and returned to the isogenic level (Fig. 8G). Lastly, the network burst duration decreased to the isogenic level (Fig. 8H). Taken together, these data indicate that L-type VGCC mediated Ca^2+^ influx drives altered neuronal network activity in P301L mutant neurons.

Since the enhanced AHNAK and tau interaction was only seen in inhibitory neurons, we next asked if mutant P301L tau expressed in inhibitory neurons is sufficient to drive neuronal network changes. To test this hypothesis, we expressed TauWT or TauP301L in either excitatory neurons or inhibitory neurons on a healthy subject background and assessed cell-type specific mutant tau expression on overall neuronal network activity (Fig. 9A). Western blots confirmed TauWT or TauP301L expression, whereas the control was mCherry empty vector (Fig. 9B). Expressing TauWT or TauP301L in excitatory neurons did not alter the spontaneous activity of overall neuronal firing rate (Fig. 9C). Upon stimulation, both excitatory expressing TauWT and TauP301L neurons showed enhanced network burst duration compared to the mCherry control, but there was no difference between TauWT and TauP301L neuronal network (Fig. 9D). Parallel experiments of expressing TauWT or TauP301L only in inhibitory neurons were also performed. An increase in overall spontaneous firing was observed in TauWT and TauP301L neurons compared to the mCherry control, although no change was observed between TauWT and TauP301L conditions (Fig. 9E). The overall burst rate increased in TauP301L neurons compared to the mCherry control too; however, no change was observed in TauWT neurons compared to the mCherry control and between TauWT and TauP301L neurons (Fig. 9F). Upon stimulation, inhibitory TauP301L expression led to longer evoked network burst duration compared to TauWT and mCherry control conditions (Fig. 9G).More spikes per burst were fired in the inhibitory TauP301L expressed condition compared to TauWT and mCherry control (Fig. 9H). Overall, TauP301L expression in inhibitory neurons showed a stronger functional influence than its expression in excitatory neurons, indicating that the presence of mutant P301L tau in inhibitory neurons is sufficient to alter neuronal and network firing patterns.

**Figure 9.**
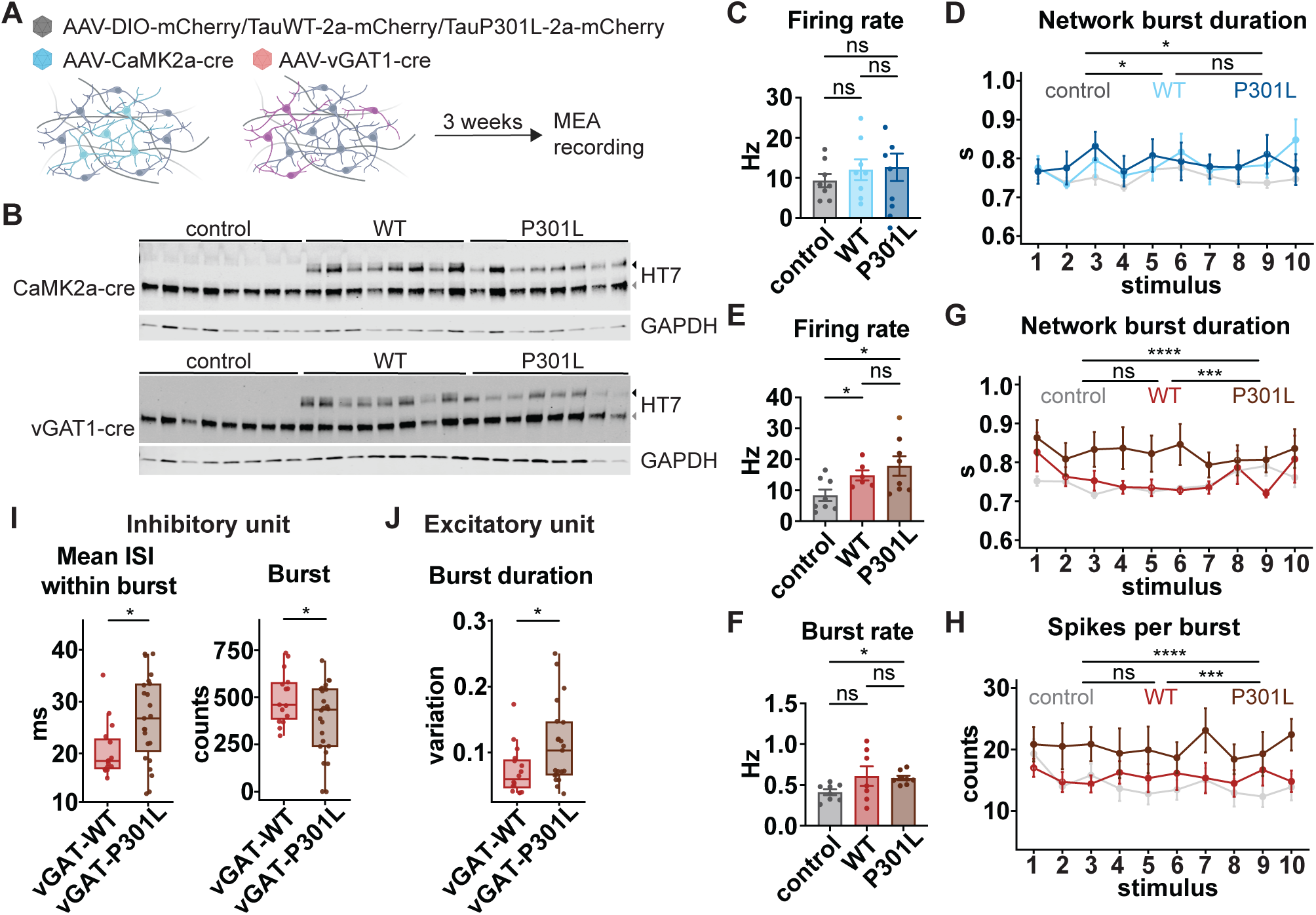
P301L tau expression in WT healthy neurons alters neuronal and network activity at both cellular and network levels. (A) Schematics of the experimental design to express wildtype hTau-2N4R (WT) or hTau carrying P301L mutation (P301L) only in excitatory or inhibitory iPSC-iNs on a healthy subject background. (B) Western blots of exogenous and endogenous tau expression in control, WT and P301L tau expressing cultures. (C-F) MEA recordings of (C) spontaneous firing rate and burst rate, (D) evoked spikes during electrical stimulations (E) ISI within network burst during stimulations, and (F) network burst duration during stimulations in neuronal cultures that selectively express control, WT and P301L in excitatory neurons. (G-J) MEA recordings of (G) spontaneous firing rate and burst rate, (H) evoked spikes during electrical stimulations (I) ISI within network burst during stimulations, and (J) network burst duration during stimulations in neuronal cultures that selectively express control, WT and P301L tau in inhibitory neurons. (K) Mean ISI within burst and burst counts of putative inhibitory units of spontaneous firing in neuronal cultures that selectively express control, WT and P301L tau in inhibitory neurons. (L) Burst duration of putative excitatory units of spontaneous firing in neuronal cultures that selectively express control, WT and P301L tau in inhibitory neurons. (C, G) One-way ANOVA followed by Tukey’s post-hoc test. (D-F, H-J) Two-way ANOVA followed by Tukey’s post-hoc test. (K-L) Unpaired t test or Wilcoxon rank-sum test with Benjamini-Hochberg adjusted p-values. Data are mean +/- SEM; ns, not significant; *p < 0.05, **p < 0.01, ***p < 0.001, ****p < 0.0001.

Next, we asked how TauP301L expressed in inhibitory neurons drives the overall neuronal and network changes. We assessed the influence of inhibitory tau on the firing patterns of putative inhibitory and excitatory neurons, by sorting the spontaneous firing spikes into single unit activity as we did in Fig.3. We separated putative excitatory units and inhibitory units from the recordings of TauP301L or TauWT expressed only in inhibitory neurons. In the presence of TauP301L, the mean ISI within burst of inhibitory units increased, and the burst of inhibitory units decreased compared to TauWT (Fig. 9I). As a result, excitatory units had larger burst duration variations in TauP301L compared TauWT (Fig. 9L), although the exogenous tau or mutant tau were not expressed in excitatory neurons. These data show similar changes as those observed in Fig. 3A, indicating that P301L tau in inhibitory neurons not only leads to firing pattern changes in inhibitory neurons but also affects the firing patterns of excitatory neurons in the same culture.

Lastly, we asked if nimodipine can reverse the altered neuronal and network patterns in cultures that TauP301L was expressed only in inhibitory neurons. Indeed, in the presence of nimodipine, spontaneous firing rate showed no difference between TauP301L and mCherry control, or TauWT and mCherry control (Fig. S6A). During stimulations, the enhanced spikes per burst in the TauP301L condition was normalized to TauWT and mCherry control levels (Fig. S6B). Additionally, evoked network burst duration was decreased in the presence of nimodipine, and the difference between TauP301L and TauWT was smaller. Although TauP301L still fired longer network bursting than the TauWT condition, no significant difference was observed between TauP301L and mCherry control (Fig. S6C). Taken together, blocking L-type VGCC largely suppressed the functional influence of TauP301L expressed in inhibitory neurons on the overall neuronal network activity.

## DISCUSSION

Our study combined electrophysiological approaches and unbiased proteomics to reveal the underlying mechanisms of neuronal network dysfunction in iPSC-iNs with the disease-causing *MAPT* P301L mutation. In the mixed excitatory and inhibitory cortical neuronal culture, the presence of P301L tau caused hyperactivity and hyperexcitability of the neuronal network. Importantly, these electrophysiological measurements could be used to build a machine learning model to categorize disease genotypes and allow to differentiate mutant P301L neurons from the isogenic control. Further mechanistic studies revealed that P301L tau led to disproportional functional changes in excitatory vs. inhibitory neurons, resulting in an imbalance of excitation and inhibition. The enhanced excitatory synaptic transmission contributed to network hyperexcitability. Furthermore, our study revealed cell-type specific protein-protein interaction networks for tau in human neurons, which led to differential pathological influence on excitatory versus inhibitory neurons in the same *MAPT* P301L mutation background. We identified increased interaction between tau P301L and AHNAK - a key driver protein in the AD protein network, which modulated Ca^2+^ signaling through L-type VGCCs in inhibitory neurons, leading to network hyperexcitability. Overall, our study uncovered underlying mechanisms of neuronal network hyperexcitability in a human iPSC derived neuronal model carrying the familial *MAPT* P301L mutation.

The electrophysiological measurements from MEA recordings provide a relative high throughput platform for functional phenotyping of iPSC-iN derived disease models [34,35]. Simultaneously recording of spiking activity at different spatial locations within the same culture allows to interrogate neuronal network activity and understand how disease-causing factors change network dynamics [34,35]. Our study revealed that *MAPT* P301L mutation caused neuronal and network hyperexcitability in human iPSC-derived neurons. Notably, the P301L mutation located in *MAPT* exon 10 is in 4R tau but absent in 3R tau. Our study and others [36] demonstrated that, despite expressing low 4R tau levels, iPSC neurons carrying this mutation still displayed tauopathy-relevant phenotypes, indicating that even low-level exon 10 inclusion impacts cellular function [36]. In our MEA functional study, the presence of P301L tau increased spontaneous spike and bursting activity, which led to increased network burst duration. The increased neuronal and network excitability also persisted during electrical stimulation, in which mutant neurons fired more irregular spikes with longer burst duration compared to the isogenic control. A previous study expressing 0N4R wildtype tau or P301L tau in mouse primary neurons observed similar phenomena [37]. Compared to wild type tau or empty vector control, increased basal spike and burst activity were observed with mutant P301L tau expression, which is consistent with our observations. Another study using rat primary cortical neurons reported different observations, in which a dual-nodal in vitro cortical neuronal network was established [38]. Expression of P301L tau in the presynaptic node reduced spike and network activity compared to the unperturbed network control. However, the duration and size of network burst, as well as network synchrony increased. In addition, the responses of presynaptic nodes were attenuated by electrical stimulation, while the postsynaptic responses were completely abolished in concert with the retraction of presynaptic axons. We observed the same increased network burst duration in iPSC-iNs carrying P301L mutation. Nevertheless, human neurons express more tau isoforms compared to mouse neurons, which likely explain the functional discrepancy in different *in vitro* cell models.

Using iPSC-derived human neurons, altered neuronal network activity was reported in other *MAPT* mutations, including V337M, S305N/S/I, and R406W. In iPSC derived cortical glutamatergic neurons, V337M did not alter spontaneous firing rate and burst rate. However, V337M mutant neurons showed increased network burst duration, spikes per network burst and network synchrony. Chronic depolarization increased firing rate in V337M but not in the isogenic control, suggesting impaired homeostatic plasticity in mutant neurons [20]. In mixed excitatory and inhibitory cortical neurons carrying S305 mutations (S305S, S305I, S305N), an increase in spontaneous firing rate was observed in mutant neurons compared to the isogenic control [39]. Although culture maturation differed between these two studies and our study (V337M at 4-5 weeks; S305 at 8 weeks; ours at 12 weeks of differentiation), the results were consistent in showing that mutant neurons exhibited increased spike rate, network burst duration, and spikes per burst indicating hyperexcitability. In a recent study using a cerebral organoid model with mixed neurons and glia, the R406W mutant caused hypoexcitability by reducing neuronal firing rate and network burst size [40], which is different from V337M, S305N and P301L mutations. These observations suggest mutation-dependent functional influence on neuronal network activity, which is likely to reflect the phenotype difference in human patients with these mutations.

Compared to studies above, our MEA study conducted a thorough analysis on neuronal and network activity patterns using quantitative metrics. These electrophysiological measures served as inputs to train a machine learning model which allowed us to functionally characterize the P301L tau altered neuronal activity patterns. Notably, network features provided spatial-temporal organization of spiking activity that is disrupted in the disease network. Combining both neuronal and network features, our model achieved high accuracy in predicting the activity patterns altered by P301L tau in comparison to its isogenic control. It also revealed features including spike timing variance, burst duration and network burst duration variation as top weighted features in the logistic regression model. A similar approach has been applied to functionally phenotype human iPSC-derived dopaminergic neurons carrying α-synuclein A53T mutation [41]. Further exploration is warranted to use the machine learning models as a reference point to assess the efficacy of various therapeutic interventions to rescue functional abnormalities in iPSC-iN tauopathy models.

In mixed neuronal models, one underlying mechanism of altered neuronal network dynamics likely converges on loss of excitatory/inhibitory balance. In our study, we isolated the activity changes of putative excitatory and inhibitory neurons by separating excitatory vs inhibitory units from MEA recordings. This allowed us to compare side by side the functional changes in different neuronal subtypes resulting from the same *MAPT* P301L mutation. One prominent observation was increased excitatory firing rate and burst duration, whereas there was decreased inhibitory burst rate in P301L mutant neurons. This points to an imbalance between excitation and inhibition in P301L mutant neurons. Increased burst duration of excitatory neurons was also observed in iPSC-derived cortical glutamatergic neurons carrying the V337M mutation [20], which was likely mediated by Ca^2+^ influx through L-type Ca^2+^ channel via the cAMP/PKA signaling pathway [42]. A152T tau also significantly increased excitatory neuronal firing frequency as indirectly measured by Ca^2+^ imaging [43], in which increased tau and Fyn kinase interaction promoted NMDA receptor mediated Ca^2+^ influx and neuronal hyperactivity [43,44]. To the best of our knowledge, no other study has directly measured the influence of *MAPT* mutation on inhibitory neurons.

Our electrophysiological studies using patch clamp techniques revealed details on how P301L tau altered firing properties of excitatory vs inhibitory neurons. Both intrinsic membrane properties and synaptic transmission in each neuronal subtype were fully characterized. In iPSC-iNs after 3 months of differentiation, the presence of P301L tau did not alter passive membrane properties of either excitatory or inhibitory neurons and these neurons all exhibited mature AP firing patterns. However, excitatory neurons exhibited a higher evoked firing rate, whereas inhibitory neurons exhibited a lower evoked firing rate with larger current injections, suggesting changes in intrinsic properties that contributed to evoked firing activity in different neuronal subtypes. A previous study showed earlier electrophysiological maturation in iPSC-derived P301L neurons and increased excitability overall compared to the healthy subject derived control [36]. Our study, by analyzing neurons expressing both 3R and 4R tau, bypassed the premature developmental stages of iPSC-iNs in which only 3R tau being expressed, and detected difference in intrinsic firing properties between excitatory vs inhibitory neurons.

Another difference between excitatory vs inhibitory neurons was in synaptic transmission. In excitatory neurons, excitatory synaptic transmission was significantly enhanced as indicated by the increased frequency of both spontaneous and miniature EPSCs, suggesting increased presynaptic release of glutamate in P301L mutant neurons. These changes were likely due to accumulation of pathological tau in vGlut1 enriched presynaptic terminals as revealed by immunostaining (Fig. 5D). Similar enhanced EPSC frequency was observed in iPSC-iNs with the 16+IVS10 mutation. Increased synaptic drive and upregulated glutamate receptor signaling via AMPA and NMDA receptor surface trafficking boosted the hyperexcitability of excitatory synaptic transmission in 16+IVS10 mutant neurons [19]. Presynaptic accumulation of pathological tau was reported in AD and FTD patient brains [45–47], and tau association with synaptic vesicles caused presynaptic dysfunction in a Drosophila tauopathy model [48]. Enhanced excitatory synaptic transmission in excitatory P301L mutant neurons was likely synergistic with their enhanced evoked excitability and contributed to hyperactivity of excitatory neurons. In iPSC-derived inhibitory neurons, we observed changes in both excitatory and inhibitory synaptic transmission. The frequency did not change, but the amplitude was significantly affected, suggesting altered postsynaptic signaling in inhibitory P301L mutant neurons. We observed opposite changes in sEPSC and mEPSC amplitude in inhibitory mutant neurons compared to the isogenic control. The increased amplitude of sEPSC was likely attributable to the hyperactivity of excitatory network. However, decreased amplitude of the mEPSC suggested homeostatic downregulation of postsynaptic excitatory signaling, which was likely synergistic with the decreased evoked excitability and contributed to the hypoactivity of inhibitory P301L mutant neurons. In addition, the decreased IPSC amplitude was likely attributable to tau accumulation in inhibitory synapses, as indicated by the colocalization between ptau and GABA positive puncta (Fig. 5C). Transcriptional deficits of GABAergic signaling pathways were reported in iPSC-iNs carrying the R406W mutation and in human brain tissues with the same mutation, in which broad downregulation of GABA receptors was observed [14]. Taken together, our data show mutant P301L tau lead to enhanced activity of excitatory neurons, but decreased activity of inhibitory neurons, which can contribute to neuronal network hyperexcitability.

Enhanced excitatory synaptic transmission through AMPAR is sufficient to drive network hyperexcitability. When the AMPAR antagonist NBQX was applied into the culture, neuronal and network hyperexcitability decreased (Fig. S5). Spontaneous firing rate and ISI variations were corrected to the isogenic level. The evoked spikes and network burst duration were also decreased. Given that the miniature EPSC only increased in excitatory neurons, we speculated that the enhanced excitatory synaptic transmission in excitatory neurons is one of the driving forces for network hyperexcitability in P301L mutant neurons. Surprisingly, blockade of NMDAR neither changed the spontaneous firing rate or pattern, nor affected evoked neuronal and network activity, suggesting that NMDAR pathways did not play a role in altered activity of iPSC-iNs with the P301L mutation. These findings differ from a previous report on the A152T tau mutation [43].

To further reveal the molecular basis that mediates pathological and functional differences in excitatory vs inhibitory neurons, we profiled cell-type specific protein-protein interaction networks for tau in human neurons. Previous tau interactome studies have reported numerous tau- or tau aggregate-associated proteins across human tissues [49], human cell models [24,50] and mouse models [51–53]. While findings in each study depended on models and tau species, these reports repeatedly emphasized tau interacting proteins associated with microtubule/cytoskeletal proteins, molecular chaperones, ribosomal proteins, RNA-binding proteins and proteostasis machinery, presynaptic vesicles and mitochondria [24,49–54]. None of these studies addressed the cell-type-specific difference of tau interactomes. Our proteomic results identified both converged and unique tau interactomes in human iPSC-derived excitatory vs inhibitory neurons. The common tau interactors in both of these neuronal subtypes were enriched in structural and cytoskeletal proteins, protein folding chaperones, cell adhesion molecules and ubiquitin protein ligase binding, which represented canonical physiological roles of tau, and were also consistent with previous reports on iPSC-derived neurons and rodent models [24,53]. Most importantly, our study also identified unique tau interactors in either excitatory or inhibitory neurons. The unique tau interactors in excitatory neurons were enriched with RNA binding proteins associated with spliceosome, ribonucleoprotein complex, and RNA metabolism. Others have shown that tau interacts with RNA-binding proteins including ribosomal proteins in human neurons [24] and in different tauopathy mouse models [51,52]. Associations of tau inclusions with RNA-binding proteins may contribute to early tau pathogenesis [51,52]. Our study also revealed unique tau interactors in inhibitory neurons, which regulate ion transporter activity and protein kinase inhibitory activity, and were enriched in dendritic spines, postsynaptic cytoskeleton and extracellular vesicles. Pre-synaptic and post-synaptic enrichment of tau interactors in human excitatory neurons has been reported in previous studies [24,55], in which tau may modulate neurotransmitter release and synaptic plasticity. The difference between the unique tau interactomes in excitatory vs. inhibitory neurons indicates different physiological roles of tau in each neuronal subtype.

Tau interactors altered by P301L tau in excitatory or inhibitory neurons reciprocally correlated with AD progression, indicating that these interactions might play important roles in disease progression. Strikingly, one of the top key drivers in the protein module most associated with AD - AHNAK, was identified as a tau interactor in inhibitory neurons which showed enhanced interaction with P301L tau. We validated this interaction and provided evidence that supported increased P301L tau/AHNAK interaction may disrupt VGCC function. As a result, increased Ca^2+^ influx in inhibitory neurons led to neuronal network hyperexcitability in neurons carrying the P301L tau mutation. Our results support an important role of AHNAK in conveying tauopathy-induced functional deficits in human inhibitory neurons. Wang et al 2025 reported that AHNAK is upregulated in AD and associated with both neuropathological traits and cognitive function [28]. Protein co-expression analyses revealed AHNAK as a hub gene in the top-ranked glia-neuron module, with highest expression in astrocytes [28]. AHNAK downregulation reduced ptau level in both human astrocytes and astrocyte/neuron coculture, which was beneficial in promoting neuronal activities [28]. That study and ours together implicate divergent roles of AHNAK in neurons and astrocytes, however, association of AHNAK with ptau is the convergent mechanism underlying pathogenesis. Further studies on how AHNAK modulates neuron-astrocyte interaction and clarification of its role *in vivo* are needed.

In summary, we have identified neuronal subtype-specific tau interactome changes that provide a molecular framework for understanding how the same *MAPT* mutation produces divergent functional consequences in excitatory vs. inhibitory neurons. In particular, the enhanced synaptic transmission resulting from ptau accumulation in excitatory synapses contributes to network hyperexcitability. In addition, the enhanced interaction between mutant tau and AHNAK in inhibitory neurons, together with the rescue effects of nimodipine, supports a model in which pathological tau disrupts calcium homeostasis in inhibitory neurons and thereby drives network-level hyperexcitability. These findings place excitatory synaptic dysfunction and inhibitory neurons at the center of early circuit dysfunction in tauopathy and suggest that subtype-specific mechanisms should be considered when developing therapeutic strategies.

## LIMITATIONS OF THE STUDY

This study uses human iPSC-derived cortical neurons to study neuronal network dysfunction. The majority of cells in the cultured condition are neurons with limited glia which can alter neuronal and network activities. Further studies use neuron-glia coculture are needed to examine the role of pathological tau in glia function and its influence on neuronal network dynamics. Immunoprecipitation followed by mass spectrometry allows detection of steady-state tau interactomes, which are proteins that exhibit a relatively strong interaction with tau. Many additional interacting proteins that weakly or transiently interact with tau are not identified in our study.

## AUTHOR CONTRIBUTIONS

C.J. and E.M.S. conceived the project and wrote the manuscript. C.J. performed the experiments and analyzed results. Y. Z. performed analyses using machine learning approaches on MEA recordings. H. E-B. performed mass spectrometry of tau interactome. S.S. provided guidance and consultation on patch clamp recording and data analyses. N.C., A. L., A. L. helped maintaining cell cultures. O. D. generated iPSC derived neural progenitor cells on the healthy subject background, T. N. provided guidance on the tau interactome study. D. S. and H. D. S provided advice on manuscript improvement. E.M.S. supervised the project.

## ACKNOWLEDGEMENT

This study was supported by NINDS and NIA R01 AG032611, R01 NS077239, R01 NS120488 (E.M.S.), Alzheimer’s Association Research Fellowships AARF-22-926735 (C.J.), and NIA K01 AG092910 (C.J.), NINDS R61NS125280, R33NS125280 (D.S.), NIH S10 RR027990 (T.A.N.), Rainwater Tau Leadership Fellowships from the Rainwater Charitable Foundation and Fellowship in Neuroscience from the Leon Levy Foundation (Y.Z.). We thank Celeste Karch (Washington University in St. Louis) for providing the F0510.2dE07 and F0510.2 iPSC lines. Generation and distribution of these resources were supported in part by the Washington University Knight Alzheimer Disease Research Center (P30AG066444), the Rainwater Charitable Foundation, and the NINDS repository. We also thank NINDS Human Cell and Data Repository for providing NN0004865 and NN0000045 iPSC lines. MEA experiments were supported by the Stem Cell Engineering Core (RRID:SCR_027503) at the Icahn School of Medicine at Mount Sinai. We thank Samuele Marro (Icahn School of Medicine at Mount Sinai) for providing instructions on using the Axion Maestro Pro MEA instrument. We also thank New York University (NYU) Langone Health’s Ion Laboratory (RRID: SCR_021754) for the technical support on electrophysiological experiments. We thank P. Davies (Albert Einstein College of Medicine and Long Island Jewish Medical Center, Litwin-Zucker Research Center at Feinstein Institutes for Medical Research) for the tau antibodies PHF1, which were obtained subsequently from P. Marambaud and J. Koppel at the Feinstein Institutes.

## Data and materials availability

All data needed to evaluate the conclusions in the paper are present in the paper and/or the Supplementary Materials. Proteomics LC-MS/MS data (raw mass spectrometry data, peak lists and results) that support the findings of this study are available at MassIVE (UCSD) (https://massive.ucsd.edu/ProteoSAFe/static/massive.jsp).

## METHODS

### Cell cultures

We obtained two lines of human induced pluripotent stem cells (iPSCs) carrying familial *MAPT* P301L mutation and their isogenic controls from the Tau consortium (Table 1) [56] and a healthy wildtype control KOLF2.1J cell line from the Jackson Laboratory [57]. The iPSCs were maintained in mTeSR™1 medium (STEMCELL Technologies, Cat# 85850) on Corning® Matrigel® hESC-Qualified Matrix-coated (Corning, Cat# 354277) tissue culture plates at 37°C and 5% CO₂. Cultures were passaged using Accutase (STEMCELL Technologies, Cat# 100-0485) and routinely assessed for pluripotency and spontaneous differentiation. Neural induction was performed using the STEMdiff™ SMADi Neural Induction Kit (STEMCELL Technologies, Cat# 08581) following the instruction to direct neuroectodermal specification through dual inhibition of TGF-β/BMP-dependent SMAD signaling. Embryoid bodies (EBs) were formed and replated onto poly-L-ornithine (PLO; 15 µg/mL in PBS; Sigma, Cat# P4957) and laminin (10 µg/mL in DMEM/F-12; Sigma, Cat# L2020) coated plates to allow neural rosette formation. Neural rosettes were selected and expanded to obtain neural progenitor cells (NPCs). NPCs were further expanded in Neural Progenitor Medium (STEMCELL Technologies, Cat# 05833), followed by either cryopreservation or further maturation in Neuron Maturation Medium (BrainPhys™ Neuronal Medium N2-A & SM1 Kit, STEMCELL Technologies, Cat# 05793). Terminal neuronal differentiation was performed using NPCs from less than 5 passages. Neurons were maintained for at least 12 weeks before assays. Half-medium changes were performed every 3 - 4 days. Mature forebrain neurons were characterized by expression of class III β-tubulin (TUJ1) to confirm neuronal identity, FOXG1 to verify forebrain cortical identity, and ability of repetitive firings of action potentials (AP). The culture consists of a mixed population of excitatory and inhibitory forebrain cortical neurons.

### Multielectrode array (MEA)

We used Axion Maestro Pro multielectrode array (Axion Biosystems). Neurons terminally differentiated in Neuron Maturation Medium for 7 weeks were replated onto a Cytoview 48-well MEA plate at a density of 100,000 cells per well. Each well contains 16 electrodes arranged in 4×4 grid. The plates were coated with PLO/laminin before seeding neurons that were dissociated by Accumax. Replated neurons were allowed to reform neural network for another 5 weeks before recording. Neuronal activity was recorded by the AxIS Navigator software (Axion Biosystems) at a sampling rate of 12.5 kHz. Spike activities were detected using a threshold set to 6 times of standard deviation of the noise. Spontaneous neuronal activity was recorded for 15 min. Evoked activity was recorded for 1 min of baseline activity followed by 10 bursts of 20 Hz biphasic pulses at 500 mV voltage. An additional 4 min of spontaneous activity was also recorded after stimulation. The Neural Metric Tool (Axion Biosystems) was then used to analyze parameters of individual spikes, bursts and network bursts. Bursts were detected at single channel level with minimum 5 spikes and maximum 100 ms inter-spike intervals. Both spike and burst activity were then analyzed at the well level. Network bursts were detected at the well level, with minimum 50 spikes, maximum 100 ms inter-spike interval, and minimum of 25% participating electrodes.

### Spike sorting

Spike sorting was performed following the Axion protocol. Specifically, the raw files of spontaneous activity were processed in AxIS Navigator and then by Axion Data Export Tool (Axion Biosystems). The converted files were further analyzed in the Offline Sorter (Plexon). Automatic sorting was first performed for all the channels using the T-Distribution E-M automatic sorting method. Manual inspection of putative units was performed according to the PCA clustering and the peaks of aligned waveforms. This step was performed in multiple iterations until a set of units that were distinct from one another based on their peak location, trough location, and waveform shape dissimilarity. Units with a firing rate less than 0.1 Hz were excluded. To identify excitatory- and inhibitory units, auto-correlogram and cross-correlogram of spikes detected on each electrode were plotted in Offline Sorter. The distribution of firing events of a test unit within a 50 ms window was centered around each spike of a reference unit. An increase in firing at small positive lags indicates that the reference unit showed excitatory effect on the test unit. A decrease in firing at small positive lags indicates that the reference unit showed inhibitory effect on the test unit. The putative excitatory- and inhibitory units detected by each electrode were saved separately. The resulting files of all excitatory units or all inhibitory units from a whole plate were then imported into the Neural Metric Tool (Axion Biosystems), followed by analyses of parameters of individual spikes, bursts and network bursts.

### Multielectrode array feature analysis and genotype classification

MEA recordings of spontaneous activity of P301L tau and isogenic neurons were analyzed at the well level. Features were organized into three electrophysiological domains: spike-level metrics, burst-level metrics, and network burst metrics. First, wells were sequentially filtered by activity thresholds (>3 active electrodes, >10 bursts, and >1 network burst) before retaining only those with complete genotype and relevant feature data for each model. Four feature groups were used for model comparison: spike features, burst features, network burst features, and a combined feature constructed by pooling all features. Genotype classification was performed using L1-regularized logistic regression. All predictors were standardized before model fitting. To compare the discriminative value of the different feature groups, we evaluated each model using stratified two-fold cross-validation and repeated the cross-validation analysis 20 times. We used a stratified 50:50 train/test split to evaluate the performance of the full combined model and generated confusion matrices for both the training and test sets. To evaluate classification performance, we performed receiver operating characteristic (ROC) analysis on the held-out test set and calculated area under the ROC curve (AUC). To assess the stability of the ROC estimate, 1,000 bootstrap iterations were generated by repeatedly drawing new stratified 50:50 train/test splits, refitting the L1-regularized logistic regression model, and recalculating AUC on each corresponding test set. The bootstrapped AUC distribution was summarized and compared with the observed AUC from the primary test split. To determine how many training wells were required for reliable genotype classification, we re-estimated model performance across a range of stratified train/test fractions from 5% to 45% training data, with 200 repeated splits per fraction. Individual features highlighted by the logistic regression model were plotted as point plots. Group differences were tested with Mood’s median test. All analyses of multielectrode array recordings were performed in Python using pandas, NumPy, SciPy, scikit-learn, statsmodels, seaborn, and matplotlib packages.

### Patch clamp recording

Neurons were grown on coverslips and infected with AAVs for 2-3 weeks before patch clamp recording. Whole-cell recording was performed on the individual neuron expressing either GCamP7s or GFP under CamK2a or vGAT driven cre recombinase. Cells were kept in the external solution containing (in mM): 140 NaCl, 2.5 KCl, 1.8 CaCl_2_, 1.2 MgCl_2_, 10 glucose, and 10 HEPES (pH 7.35, 295-300 mOsm). Patch pipettes were pulled from borosilicate glass (World Precision Instruments, 140N-15) with an access resistance of 3–5 MΩ. These pipettes were filled with an internal solution containing (in mM): 135 K-gluconate, 8 KCl, 10 phosphocreatine, 10 HEPES, 3 Mg-ATP, and 0.3 Na-GTP, 2 MgCl_2_ (pH 7.25, 277 mOsm). After membrane breaking to form whole cell configuration, the passive membrane characteristics including membrane resistance, cell capacitance, and resting membrane potential were measured for each neuron. Subsequently, a series of depolarization current steps were injected from a holding potential of −70 mV to elicit action potentials (APs). Data were recorded at a sampling rate of 10 kHz and filtered at 2 Hz by an 8-pole Bessel filter using a MultiClamp 200 amplifier (Molecular Devices). Cells with resting membrane potential more depolarized than −50 mV and access resistance exceeding 25 MΩ were excluded from further analyses. AP frequency as a function of injected currents were analyzed by custom-written Matlab scripts.

To record spontaneous excitatory and inhibitory postsynaptic currents, the internal solution to record AP was used. The membrane potential was held at −70 mV to record excitatory postsynaptic currents (EPSCs), which was at the reversal potential of Cl^−^. Then the membrane potential was slowly increased and held at 0 mV, which was at the reversal potential of cations, to record inhibitory postsynaptic currents (IPSCs) [58]. To record miniature postsynaptic currents, 1 μΜ tetrodotoxin (TTX) was added into the external solution. The internal solution was switched to Cs-based solution containing (in mM): 130 Cs-methanesulfonate, 4 TEA-Cl, 5 CsCl, 1 QX-314, 10 phosphocreatine, 3 MgATP, 0.3 NaGTP, 10 HEPES, and 1 EGTA for EPSCs. Data were analyzed in Clampfit (Molecular Devices).

### Western blotting

Neurons were rinsed in DPBS and then homogenized in ice-cold radioimmunoprecipitation assay (RIPA) buffer (50 mM Tris-HCl, 150 mM NaCl, 1 mM EDTA, 1% Nonidet P-40, 0.25% sodium deoxycholate, 0.1% SDS, pH 7.4) with a protease cocktail (Roche) and a phosphatase inhibitor cocktail (Sigma). Cell homogenates were collected and incubated on ice for 15 min before being centrifuged at 14,000 x g for 15 min at 4 °C. The resulting supernatants were collected, and concentrations were assessed using a BCA assay (Thermo Fisher Scientific). Equal amount of protein lysates was eluted in sample buffer, and boiled for 10 min, followed by electrophoresis on SDS-polyacrylamide gel electrophoresis (PAGE) gel and transferred onto nitrocellulose membrane.

Membranes were blocked by 5% milk in Tris buffered saline with 1% TritonX-100 (TBST). Primary antibodies, including NeuN (1:1000), GAD1 (1:1000), CaMK2a (1:1000), PHF1 (1:5000), pT231 (1:2000), AT8 (1:1000), HT7 (1:1000), Tau5 (1:1000), GAPDH (1:5000), TUBB (1:5000), 3R-tau (1:1000), and 4R-tau (1:1000) were diluted in Superblock (Thermo Fisher Scientific), and incubated with membranes at 4 °C overnight. The following day the membrane was washed 4 x 10 min, and incubated with diluted (1:10,000) IRDye 800CW goat anti-rabbit or IRDye 680RD goat anti-mouse secondary antibodies (LICOR Biosciences). After another 4 x 10 min wash, blots were then detected by Odyssey CLx imaging system (LICOR Biosciences).

### Immunofluorescence

Cells growing on coverslips were washed in HBSS, followed by fixation in 4% paraformaldehyde and permeabilization in 0.2% Triton-X-100 at room temperature for 15 min of each. Fixed and permeabilized cells were blocked in 5% bovine serum albumin and 5% normal donkey serum at room temperature for 1 h. Subsequently, cells were incubated with diluted primary antibodies in HBSS with 5% bovine serum albumin and 1% normal donkey serum at 4 °C overnight. The dilution of the primary antibodies was as follows: GABA (1:500), MAP2 (1:1000), vGlut1 (1:100), TUBB (1:1000), and AT8 (1:100). On the following day, the cells were wash with HBSS 3 x 5 min and then incubated with diluted secondary antibodies (1:1000) at room temperature for 1 h. After washing with HBSS 4 x 10 min, coverslips were mounted in ProLong Diamond antifade reagent (Thermo Fisher Scientific, P10144). Images were taken by an LSM 800 Zeiss confocal laser scanning microscope and processed by Fiji ImageJ software.

### Flag pulldown of tau

Neurons were infected with AAV-CaMK2a-cre or AAV-vGAT-cre with AAV-CAG-DIO-Flag-TauWT or AAV-CAG-DIO-Flag-TauP301L to have TauWT or TauP301L expressed in either excitatory neurons or inhibitory neurons (CaMK2a-TauWT, CaMK2a-TauP301L, vGAT-TauWT, vGAT-TauP301L). After 3 weeks of AAV infection, neurons were washed with DPBS once followed by homogenized in cold lysis buffer (50mM Tris, 150 mM NaCl, 1 mM EDTA, 0.5% NP40, pH 7.4) with protease inhibitor (Roche) and phosphatase inhibitor cocktails (Sigma). The lysates were subjected to 3 repetitive freeze/thaw cycles and then centrifuged at 14000 x g for 20 min at 4 °C. Protein concentration was determined by BCA assays. Magnetic anti-Flag beads (Thermo Fisher Scientific) were washed twice in lysis buffer. Lysates were added to the beads and incubated with rotation at 4 °C overnight. The beads were then washed by lysis buffer twice and PBS twice. Bead-bound proteins were eluted by adding 3x Flag peptide (1.5 mg/mL, Thermo Fisher Scientific) in PBS for 10 min at room temperature. The elution was repeated once, and the two supernatants were combined, followed by mass spectrometry for protein identification.

### Mass spectrometry

Protein identification and quantitation were performed by liquid chromatography–mass spectrometry (LC–MS/MS) using label-free data-independent acquisition (DIA) [59]. Flag antibody enriched elution fractions were concentrated by a brief SDS-PAGE run. Proteins were visualized and fixed by staining with EZ-Run Protein staining solution (Thermo Fisher Scientific). The stained protein gel regions were excised and completely de-stained. Finally, each gel sample was dehydrated with 100% acetonitrile. Cysteinylated proteins were reduced with 10 mM dithiothreitol and alkylated with 55 mM iodoacetamide. After two washes with 50 mM NH_4_HCO_3_ in 30% acetonitrile, samples were dried by vacuum centrifugation. Dried gel samples were digested overnight with 500 ng mass spectrometry-grade trypsin (Trypsin Gold, Promega, Madison, WI, USA) in 50 mM NH_4_HCO_3_ digestion buffer. After acidification with formic acid, peptides were extracted with 5% formic acid/50% acetonitrile (v/v) and concentrated to a small droplet using vacuum centrifugation. Desalting of peptides was performed using hand-packed SPE Empore C18 Extraction Disks as described [60]. Desalted peptides were again concentrated and reconstituted in 10 μL of 0.1% formic acid in water. Aliquots of the peptides were analyzed by nano-LC-MS/MS using an Easy nLC 1000 equipped with a self-packed 75 μm × 20 cm reverse phase column (ReproSil-Pur C18-AQ, 3 μm, Dr. Maisch GmbH, Germany) coupled online to a Q Exactive HF Orbitrap mass spectrometer via a Nanospray Flex source (all instruments were from Thermo Fisher Scientific, Waltham, MA, USA). Analytical column temperature was maintained at 50°C by a column oven (Sonation GmbH, Germany). Peptides were eluted with a 3%–40% acetonitrile gradient over 110 min at a flow rate of 250 nL/min. The mass spectrometer was operated in DIA mode. MS survey scans were acquired in profile mode at a resolution of 120,000 (at m/z 200) over a scan range of 300–1650 m/z. Following the survey scans, 30 groups of precursors were selected for fragmentation with sliding isolation windows to include peptide m/z values ranging from 364 to 1370 Th. The default maximum charge state was set to 4, and resolution was set to 30,000. In MS/MS, the fixed first mass was set to 200 Da. The normalized collision energy (NCE) / stepped NCE was 25.5, 27, and 30. The maximum injection time for the survey scan was 60 ms, and for MS/MS, it was set to auto. The ion target value for both scan modes was set to 3 × 10^6^. Data were analyzed by MaxQuant software (version 2.5.1.0), termed the MaxDIA analysis type.

To identify peptides, we used human in silico-generated spectral libraries that were created from the UniProt human protein sequence database (downloaded on 11/14/2025; 13,821 entries plus additional fasta of isoforms) from the Max Planck Institute of Biochemistry data-sharing drive (MPIB Datashare https://Datashare.biochem.mpg.de/s/qe1IqcKbz2j2Ruf?path=%2FDiscoveryLibraries). To determine the relative abundance of a protein across a set of samples, we used its label-free quantitation (MaxLFQ) intensity calculated by MaxDIA [61], which normalizes the protein intensity based on peptide intensities across samples.

### Mass spectrometry data analyses

We compared gene-level log_2_-transformed intensity values between experimental groups (CaMK2a-TauWT, CaMK2a-TauP301L, vGAT-TauWT, vGAT-TauP301L, n=4 replicates per group). To identify tau interactors by Flag pulldown, proteins detected in more than 3/4 of replicates for each condition were included for further analyses (Table 3). To determine proteins that had altered interaction with TauWT or TauP301L (Table 4), proteins were required to meet one of two following criteria as in Tracy et al. 2022 [24]: 1) the protein presented in both groups and significantly different with adjusted p value < 0.05, or 2) the protein detected in more than 3 replicates in one group but less than 1 replicate in the other group. Enriched or depleted proteins were subjected to either STRING Gene Ontology functional enrichment analysis [62], or ClueGO version 2.5.10 in Cytoscape version 3.10.0 [63,64]. Three ontologies were used, including GO Molecular Function, GO Biological Process, and GO Cellular Compartments.

### Analyses of P301L tau altered tau interactors in neurodegenerative disease brain

Eigenprotein computation and cross-disease comparison were performed using per-sample protein abundance values covering 802 frontal cortex samples that were extracted across eight diagnostic groups (Control, MCI, AD, FTLD-TDP, LBD, PD, PSP, VaD) from Shrestha et al. 2026 [30]. For each protein list (excitatory, 20 proteins; inhibitory, 29 proteins), a list-level eigenprotein was computed as the first principal component (PC1) of the z-scored, log-transformed abundance matrix restricted to list members present in each cohort, following the eigenprotein approach described by Tracy et al. 2022 [24]. Pairwise comparisons between each disease group and the combined control group were performed using the Wilcoxon rank-sum test with Benjamini-Hochberg false discovery rate (FDR) correction across the seven disease comparisons within each list.

Eigenprotein trajectory across AD diagnosis groups were performed using per-sample TMT protein abundance from consensus DLPFC (BA9) proteomics dataset (488 samples: 106 Control, 200 AsymAD, 182 AD; ROSMAP and Banner cohorts combined after pathology-based harmonization) from Johnson et al. 2022 [29]. List-level eigenproteins were computed as described above. Eigenprotein values were compared across the three diagnosis groups (Control, AsymAD, AD) using the Kruskal-Wallis test with post-hoc pairwise Benjamini-Hochberg correction. Diagnosis groups were defined by the original study’s pathology-based classification scheme (CERAD, Braak, and cognitive status).

Weighted cognitive resilience enrichment used per-protein effect sizes and p-values of the proteome-wide association study (PWAS) of longitudinal cognitive trajectory in the ROSMAP cohort from Wingo et al. 2019 [26]. A weighted enrichment score for each list was computed. The score was defined as the mean per-protein effect on cognitive slope weighted by −log10(p), where negative values indicate enrichment in the anti-resilience direction (higher brain abundance associated with faster cognitive decline relative to neuropathology burden). A permutation-based null distribution was generated by sampling 10,000 random protein sets of equal size from the PWAS background, and a z-score and two-tailed p-value were computed for both the pro-resilience and anti-resilience directions. Individual list proteins were classified as pro-resilience or anti-resilience based on the published FDR-thresholded PWAS categories, and enrichment within each category was tested using Fisher’s exact test with Benjamini-Hochberg correction.

KDP enrichment and degree analysis used the Bayesian causal network that was constructed from 7,269 proteins and 10,369 directed edges of parahippocampal gyrus proteomics by integrating protein quantitative trait loci, causal inference testing, and Markov chain Monte Carlo simulation with ENCODE transcription factor priors in Wang et al. 2025 [28]. Key driver protein (KDP) rankings were used to test whether each list was enriched among the 580 KDPs by Fisher’s exact test, with the background defined as all proteins present in the network. Per-protein in-degree, out-degree, and total degree were computed using the igraph R package (version 1.3.5). List-level degree distributions were compared to the full network background using two-sided Wilcoxon rank-sum tests. For visualization, 500 background proteins were randomly sampled for the violin plot.

All analyses were performed in R version 4.4.2 and packages including dplyr, tidyr, ggplot2, ggrepel, igraph for network analysis, and pwr for power calculations.

### Proximity ligation assay

Neurons were infected with AAV-hSyn-DIO-GFP and AAV-vGAT-cre for 3 weeks, followed by fixation in 4% paraformaldehyde and permeabilization in 0.2% Triton-X-100 at room temperature for 15 min of each. Cells were then washed by DPBS for 3 x 5 min. Proximity ligation assays were performed using the DuoLink In Situ Fluorescence kit (Sigma). Fixed and permeabilized neurons were incubated in a humidified plate with Duolink Blocking Solution at 37°C for 1 h. The primary antibodies (HT7, AHNAK, CACNB2, 1:200) were then diluted in Antibody Diluent and incubated with cells at 4°C for overnight. On the following day, cells were washed with Wash Buffer A for 2 x 5 min, then incubated with Duolink In Situ PLA Probe Anti-Rabbit PLUS and Duolink In Situ PLA Probe Anti-Mouse MINUS PLA probes (Sigma) at 37°C for 1 h. Cells were washed for 3 x 5 min with Wash Buffer A, followed by probe ligation at 37°C for 30 min. After two additional washes with Wash Buffer A, probe amplification was performed at 37°C for 100 min. The cells were then washed with Wash Buffer B for 2 x 10 min and mounted onto Duolink mounting media with DAPI. Images were taken under an LSM 900 Zeiss confocal laser scanning microscope and processed by Fiji ImageJ software.

### Calcium imaging

Neurons growing on coverslips were infected with AAV-hSyn-DIO-GCamP7s and AAV-vGAT-cre for 3 weeks. On the day of imaging, the coverslip was transferred into a glass bottom chamber (Warner Instruments) containing the patch clamp external solution at room temperature. Fluorescence time-lapse images were collected under an Olympus BX51WI upright microscope using a 40x objective, a fluorescent Xenon LED light source and QICAM CCD camera. Exposure time was set to 100-150 ms, and frame rate was set to 2.5 Hz. A constant flow of external solution was perfused by an AutoMate Scientific ValveLink8® Perfusion system at a rate of 1 ml per min. For each imaging session, a spontaneous activity baseline was recorded for 30 sec, followed by 30 mM KCl perfusion solution containing (in mM): 112.5 NaCl, 30 KCl, 1.8 CaCl_2_, 1.2 MgCl_2_, 10 glucose, and 10 HEPES (pH 7.35, 295-300 mOsm) for 20 sec and then 7 min KCl washout period. Images were saved as tiff files.

### Calcium imaging analyses

Calcium images were processed in Fiji imageJ. We selected ROIs covering neuronal soma. Fluorescence intensity over the time-course was measured by averaging all pixels within individual ROIs. For each KCl stimulation experiment, the baseline fluorescence intensity was averaged from 10 sec before the onset of KCl stimulation and denoted as F_0_. Upon KCl stimulation, Ca^2+^ influx was measured by relative fluorescence changes using (F - F_0_)/ F_0_, in which F is the raw fluorescence intensity at each time point. The relative fluorescence time courses (ΔF/ F_0_) were then aligned by the onset of KCl stimulation across all neurons. The average trace with SEM of all isogenic were compared to that of P301L tau neurons.

### Statistical analysis

Results from MEA, patch-clamp, imaging and western blot experiments are mean ± SEM unless stated in the figure legends. Figure 3B-F, 3H-J, 7C-D are boxplots, where the center represents median with the 25^th^ and 75^th^ percentiles marked by the box limits. The bar extends to 1/5 times the interquartile range from the median. Statistical analyses were performed in Graphpad Prism 11, R/RStudio, or Python, including unpaired t-test, Mann Whitney test, one-way ANOVA with post-hoc analyses, two-way ANOVA with post-hoc analyses, and linear mixed-effects model with post-hoc analyses. P-value < 0.05 was considered statistically significant, or as stated in the manuscript or figure legends.

**Table 1. iPSC line information.** Two pairs of patient-derived iPSC carrying *MAPT* P301L mutation and the isogenic control were used in this study. The information on the source, name and genotype of these iPSCs lines, as well as donor’s age, sex and clinical status are listed. A, asymptomatic.

**Table 2. Electrophysiological parameters measured in iPSC-iNs with tau P301L mutation and its isogenic control.** Passive membrane properties including input resistance, capacitance, and resting membrane potential (Vrest) were measured by patch-clamp experiments for each line.

**Table 3. Comparisons of tau interactors between excitatory and inhibitory neurons.** TauWT interactors in excitatory (Ex) and inhibitory (IN) neurons were listed. The top 10 Gene Ontologies in biological processes, molecular functions and cellular components of the Ex and In common interactors, Ex or In unique interactors were analyzed.

**Table 4. Comparisons of TauWT and TauP301L interactors in excitatory and inhibitory neurons.** Lists of tau interacting proteins that showed enhanced or reduced interaction with P301L tau compared to the wildtype tau.

---

**Supplemental Figure 1.**
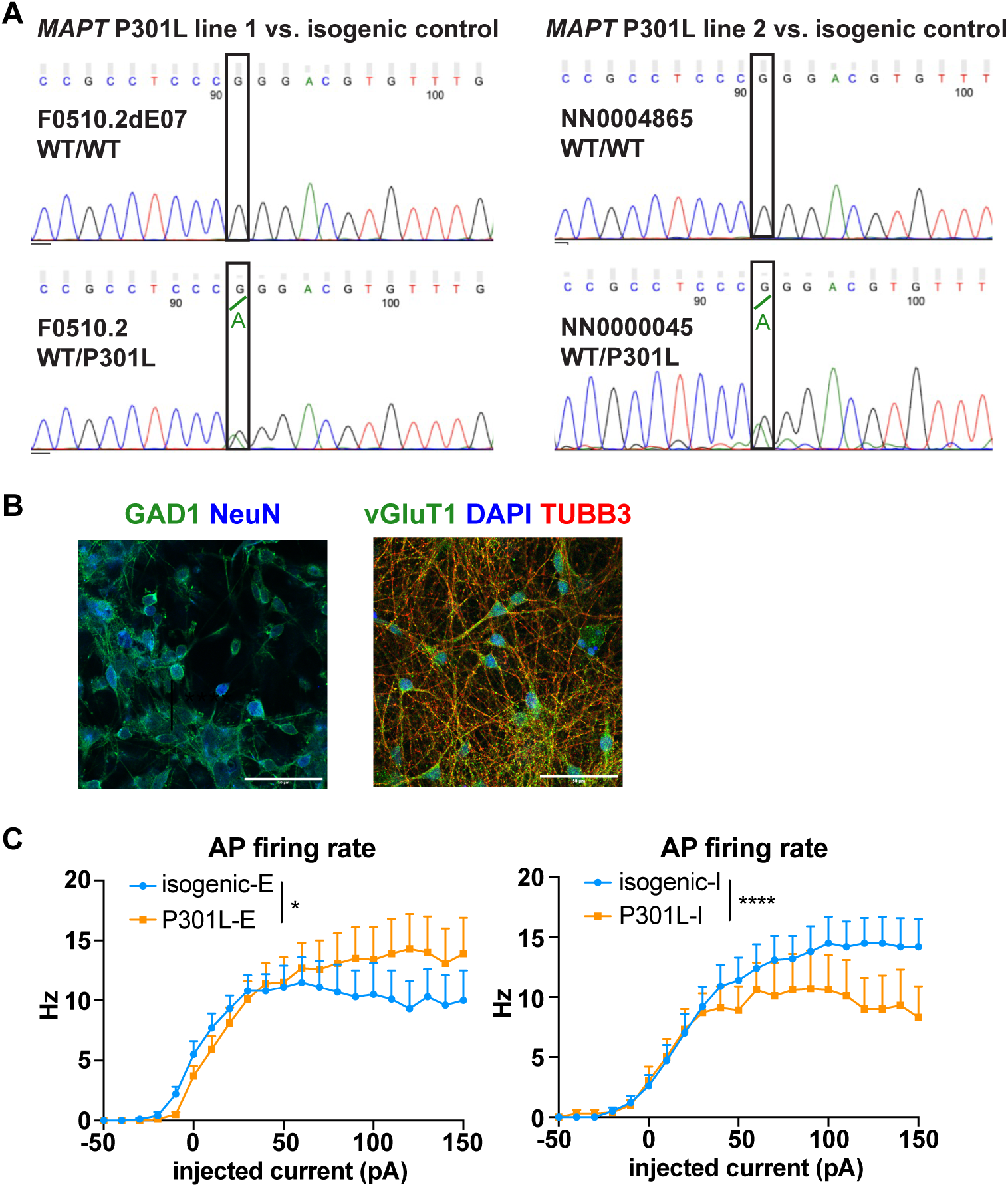
A**d**ditional **characterizations of iPSCs and iPSC-iNs carrying *MAPT* P301L mutation.** (A) Sanger sequencing chromatograms at the *MAPT* P301L site confirming genotypes of the two patient/isogenic pairs: line 1, F0510.2d2E07 (WT/WT) and F0510.2 (WT/P301L); line 2, NN0004865 (WT/WT) and NN0000045 (WT/P301L). (B) Representative images of iPSC-iNs expressing the inhibitory marker GAD1 and neuronal marker NeuN (left) and the excitatory neuronal marker vGluT1, neuronal marker TUBB3 and nuclear marker DAPI (right). (C) F/I curves of excitability in the second line of isogenic control versus P301L excitatory neurons (left) and inhibitory neurons (right). Two-way ANOVA. Data are mean +/- SEM; *, p < 0.05, ****, p < 0.0001.

**Supplemental Figure 2.**
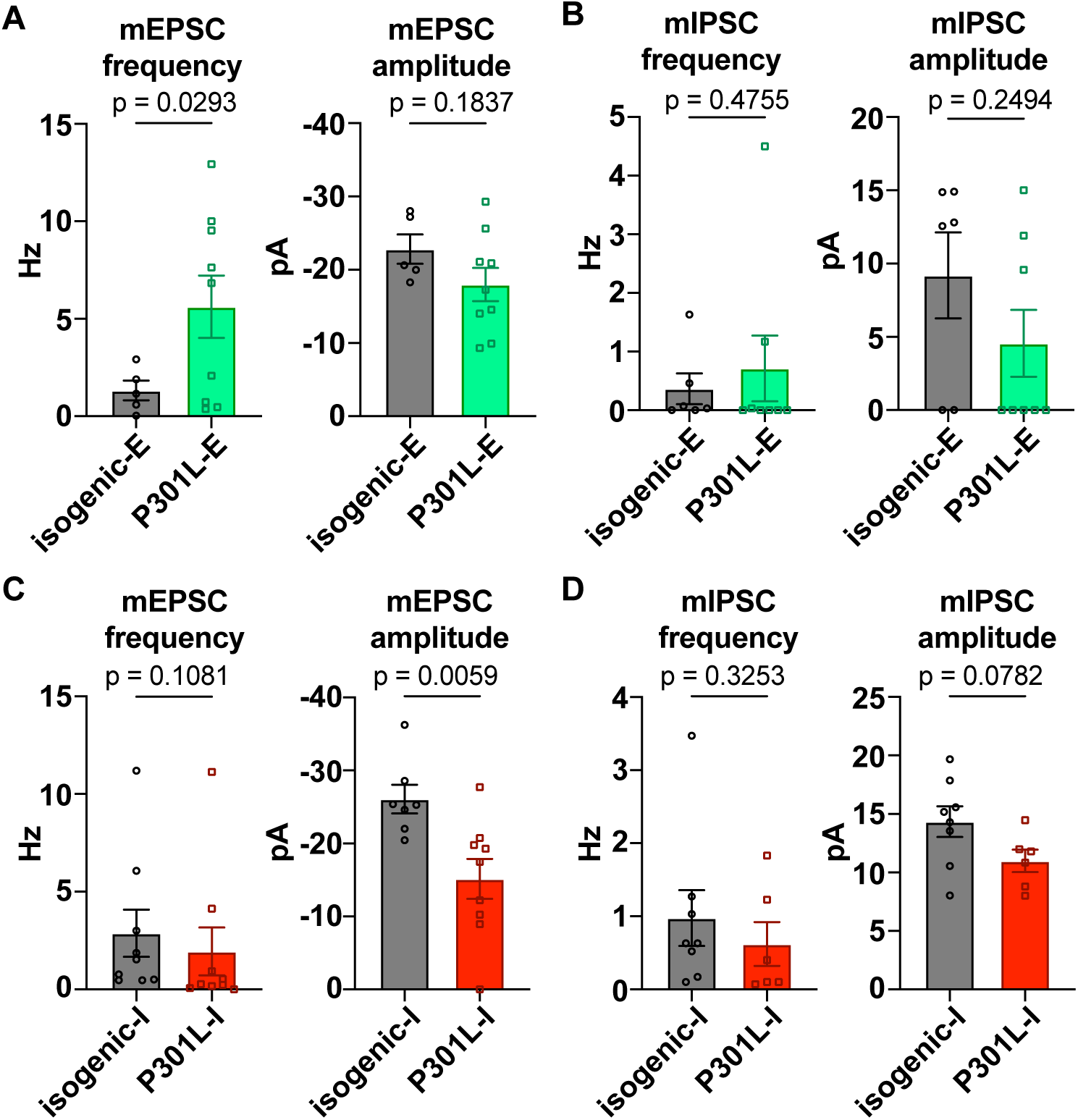
M**i**niature **post-synaptic current recording in iPSC-iNs carrying the *MAPT* P301L mutation.** Miniature post-synaptic currents (mPSC) were recorded in the presence of 1 μΜ tetrodotoxin. (A) The frequency but not amplitude of mEPSCs increased in excitatory P301L neurons compared to isogenic control. (B) Neither frequency nor amplitude of mIPSC changed in excitatory P301L neurons compared to isogenic control. (C) The amplitude but not frequency of mEPSCs decreased in inhibitory P301L neurons compared to isogenic control. (D) Neither the frequency nor amplitude of mIPSCs changed in inhibitory P301L neurons compared to isogenic control. Unpaired t test or Mann Whitney U test. Data are mean +/- SEM; p values as indicated.

**Supplemental Figure 3.**
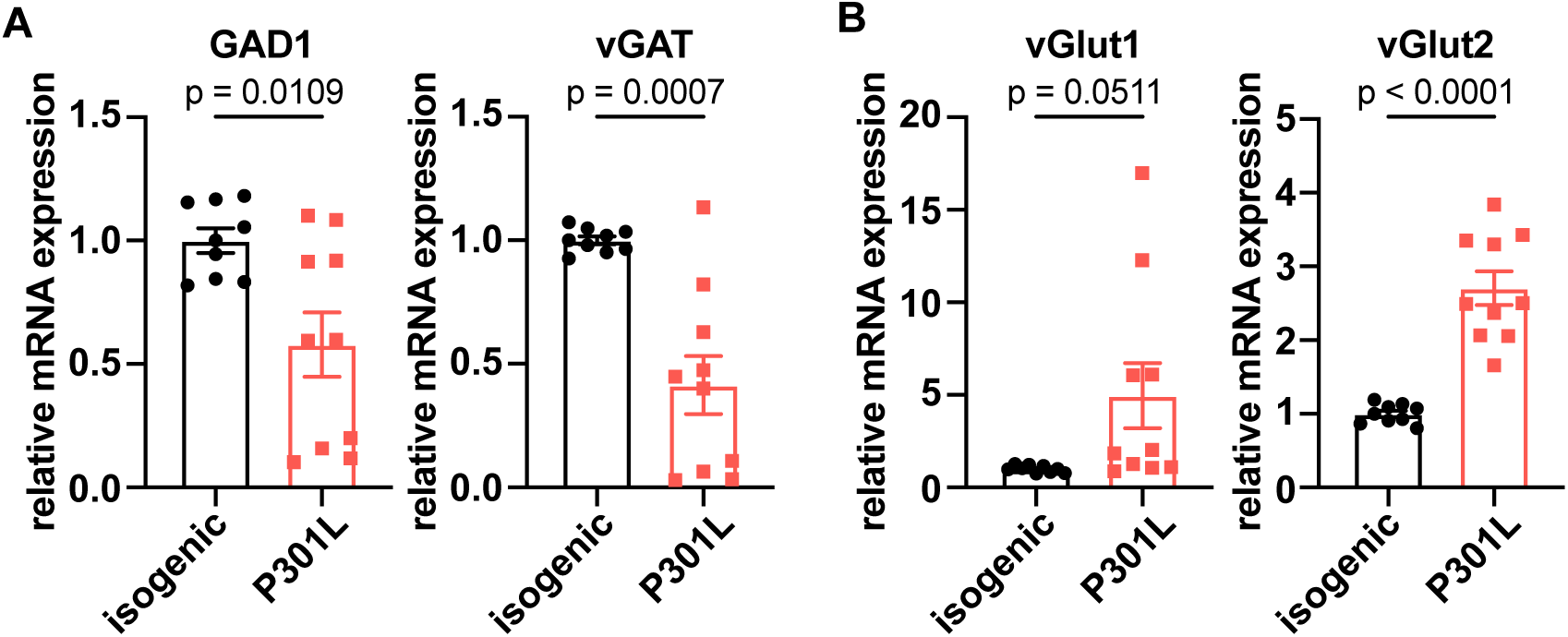
T**r**anscriptional **changes in inhibitory and excitatory markers in iPSC-iNs carrying the *MAPT* P301L mutation.** (A) Decreased expression of inhibitory neuronal markers but (B) increased excitatory neuronal markers at the transcriptional level. GAD1: GAD67, Glutamate Decarboxylase 1; vGAT: vesicular GABA transporter; vGlut: vesicular glutamate transporter. Relative mRNA expression was normalized to the expression of TUJ1, the neuron specific β-3 tubulin and compared to that in isogenic controls (ΔΔCt). Unpaired t test. Data are mean +/- SEM; p value as indicated.

**Supplemental Figure 4.**
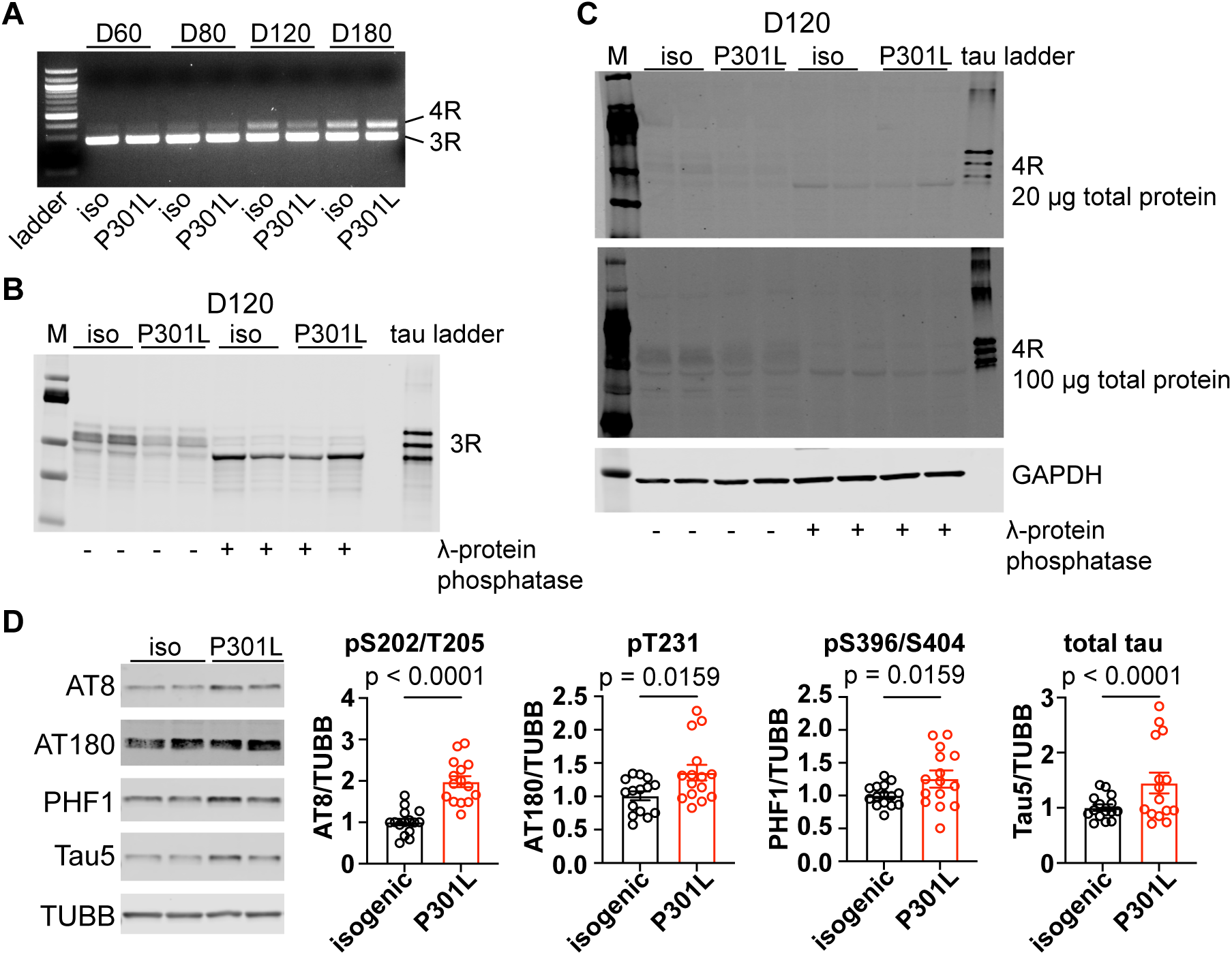
E**x**pression **of tau isoforms and phosphorylated tau in iPSC-iNs carrying the *MAPT* P301L mutation.** (A) RT-PCR of 3R and 4R tau isoforms in isogenic control (iso) and P301L iPSC-iNs at days 60, 80, 120 and 180 in vitro. (B) Western blot of 3R tau isoforms with and without λ-protein phosphatase treatment, alongside a recombinant tau ladder, showing the 3R band. (C) Western blots of 4R tau loaded at 20 μg (top) and 100 μg (middle) total protein with and without dephosphorylation; GAPDH is a loading control. (D) Immunoblotting of total tau (Tau5) and phosphorylated tau (pT231, pS202/T205, pS396/S404). TUBB, β-3 tubulin is a loading control. Quantifications shown below of the blots are from four independent batches of iPSC-iN differentiation. Relative expression was normalized to TUBB and compared to the isogenic control. Unpaired t test. Data are mean +/- SEM; p values as indicated.

**Supplemental Figure 5.**
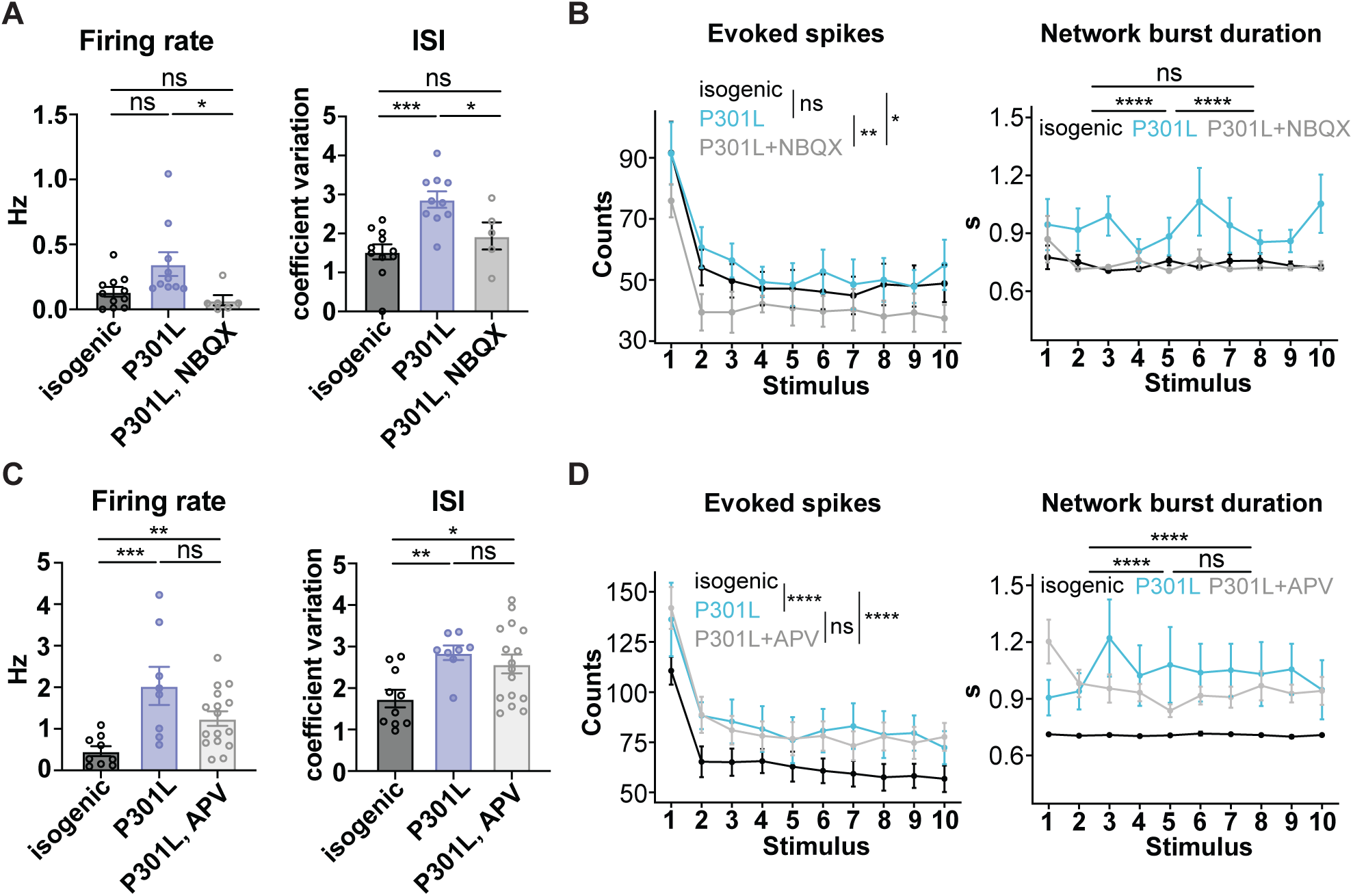
R**o**les **of AMPAR or NMDAR mediated excitatory synaptic transmission in neuronal network hyperexcitability.** (A) ΜΕΑ recordings of spontaneous firing rate and ISI coefficient of variation of spikes in isogenic control, P301L, and P301L cultures treated with the AMPAR antagonist NBQX. (B) Evoked spikes and network burst duration during electrical stimulations for isogenic control, P301L and P301L + NBQX cultures. (C) ΜΕΑ recordings of spontaneous firing rate and ISI coefficient of variation of spikes in isogenic control, P301L, and P301L cultures treated with the NMDAR antagonist APV. (D) Evoked spikes and network burst duration during electrical stimulations for isogenic control, P301L and P301L + APV cultures. (A, C) One-way ANOVA followed by Tukey’s post-hoc test. (B, D) Linear mixed-effects model followed by Tukey’s post-hoc test. Data are mean +/- SEM; ns, not significant, *p < 0.05, **p < 0.01, ***p < 0.001, ****p < 0.0001.

**Supplemental Figure 6.**
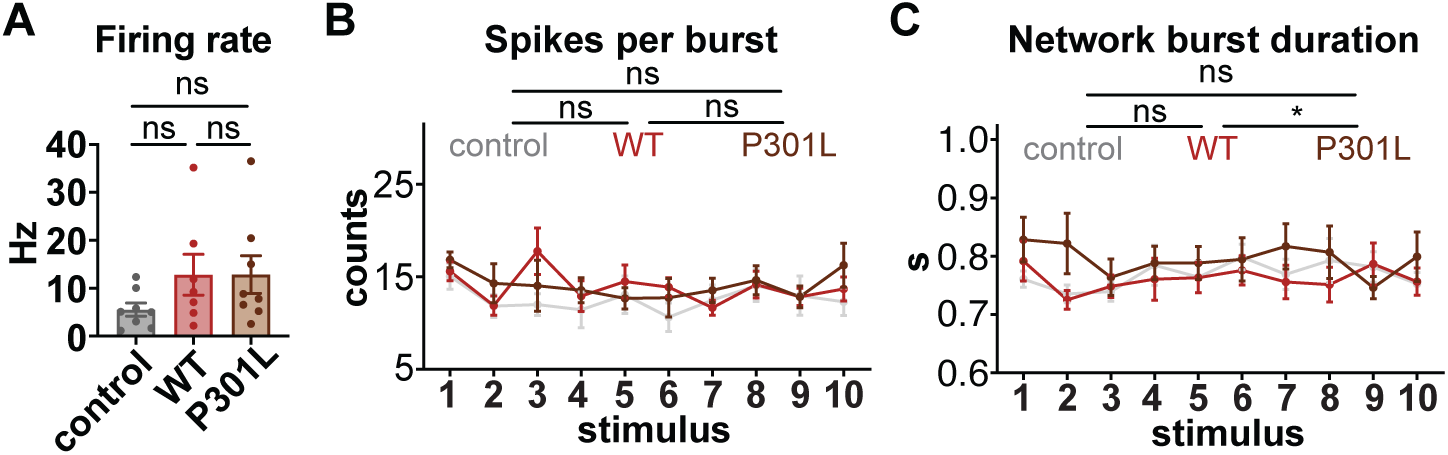
Nimodipine blocks the influence of P301L tau on neuronal and network activity. (A) Spontaneous firing rate in neuronal cultures that selectively express empty vector control, WT and P301L tau in inhibitory neurons in the presence of nimodipine. (B) Evoked spikes, spikes per burst and network burst duration during electrical stimulations in nimodipine-treated neuronal cultures that selectively express control, WT and P301L tau in inhibitory neurons. (C) Evoked spikes and network burst duration during electrical stimulations in nimodipine-treated neuronal cultures that selectively express control, WT and P301L tau in excitatory neurons. (A) One-way ANOVA followed by Tukey’s post-hoc test. (B-C) Two-way ANOVA followed by Tukey’s post-hoc test. Data are mean +/- SEM; ns, not significant, *p < 0.05, **p < 0.01, ****p < 0.0001.

